# Identification and functional characterization of transcriptional activators in human cells

**DOI:** 10.1101/2021.07.30.454360

**Authors:** Nader Alerasool, Zhen-Yuan Lin, Anne-Claude Gingras, Mikko Taipale

**Affiliations:** Department of Molecular Genetics, University of Toronto, Toronto, ON M5S 1A8, Canada; Donnelly Centre for Cellular and Biomolecular Research, University of Toronto, Toronto, ON M5S 3E1, Canada; Lunenfeld-Tanenbaum Research Institute, Mount Sinai Hospital, Sinai Health System, Toronto, ON M5G 1X5, Canada

## Abstract

Transcription is orchestrated by thousands of transcription factors and chromatin-associated proteins, but how these are causally connected to transcriptional activation or repression is poorly understood. Here, we conduct an unbiased proteome-scale screen to systematically uncover human proteins that activate transcription in a natural chromatin context. We also identify potent transactivation domains among the hits. By combining interaction proteomics and chemical inhibitors, we delineate the preference of both known and novel transcriptional activators for specific co-activators, highlighting how even closely related TFs can function via distinct co-factors. Finally, we show that many novel activators are partners in fusion events in tumors and functionally characterize a myofibroma-associated fusion between SRF and C3orf62, a potent activator. SRF-C3orf62 activates transcription in a CBP/p300-dependent manner and promotes proliferative and myogenic transcriptional programs. Our work provides a functional catalogue of potent transactivators in the human proteome and a platform for discovering transcriptional regulators at genome scale.

## INTRODUCTION

Transcription of protein-coding genes is orchestrated by a coordinated interplay of transcription factors (TFs) that bind DNA in a sequence-specific manner, RNA polymerase II machinery that initiates transcription from promoters, and diverse chromatin-associated factors and complexes that modulate chromatin structure and act as bridges between TFs and RNA pol II (Cramer, 2019). The human genome encodes thousands of proteins that are involved in various stages of transcriptional regulation, and the ready availability of methods such as ChIP-seq has revealed the genomic binding sites of hundreds of factors in diverse conditions (The ENCODE Project Consortium et al., 2020). At the same time, systematic studies have characterized or inferred the DNA-binding specificity of about three-quarters of human TFs (Badis et al., 2009; Jolma et al., 2013; Lambert et al., 2018; Najafabadi et al., 2015; Weirauch et al., 2014). Similarly, interaction proteomics approaches have uncovered many chromatin-associated proteins and characterized the composition of transcriptional regulatory complexes in human cells (Gao et al., 2012; Huttlin et al., 2020; Lambert et al., 2019; Li et al., 2015; Marcon et al., 2014; Mashtalir et al., 2018).

However, whether and how TFs and chromatin-associated factors promote transcriptional activation or repression (or regulate chromatin states by other means) has remained largely unknown due to the limited causal insights afforded by these methods. For example, the vast majority of genomic binding sites observed in ChIP-seq experiments are not causally associated with transcriptional events. That is, knockdown or knockout of a given transcriptional regulator does not affect the transcription of most genes that the regulator binds to. On the other hand, sequence-based annotation of transcriptional regulators has been challenging, because most transcriptional effector functions are encoded by degenerate linear motifs rather than folded and conserved protein domains (Arnold et al., 2018; Erijman et al., 2020; Sigler, 1988; Staller et al., 2021).

Artificial recruitment, also known as activator bypass, is powerful method to characterize the transcriptional effect of diverse proteins in a defined context (Ptashne and Gann, 1997; Sadowski et al., 1988). In this approach, proteins or their fragments are ectopically recruited to a reporter gene by fusing the protein to the DNA-binding domain of a well-characterized TF such as Gal4 or TetR. The defined context alleviates the challenges posed by endogenous gene regulation, where multiple factors bind regulatory elements in concert, hindering causal inference. Artificial recruitment has been traditionally used to identify transcriptional activators or transactivation domains (TADs) in individual transcriptional regulators (Ptashne and Gann, 1997). However, recent studies have characterized the transcriptional effects of large collections of regulators in fruit flies or yeast by individually tethering them to reporter genes (Keung et al., 2014; Stampfel et al., 2015). Due to the limited scalability of the arrayed format, these studies focused on known regulators rather than potentially novel factors. Moreover, classical model organisms lack several regulatory mechanisms and layers critical for gene expression in mammals, such as enhancers (yeast) or DNA methylation (yeast and fruit flies). More recently, Tycko and colleagues implemented an unbiased pooled screening strategy to characterize the transcriptional activation potential of annotated human protein domains (Tycko et al., 2020). This study highlighted the value of unbiased approaches in identifying novel transcriptional regulators, such as the unexpected role of variant KRAB domains in transcriptional activation instead of repression. Yet, because most transcriptional activation domains are encoded by disordered regions (Arnold et al., 2018; Dyson and Wright, 2005), it is likely that a domain-focused screen misses a significant fraction of transactivators.

Thus, despite our increased knowledge of the composition of transcriptional regulator complexes and their genomic binding patterns, we still lack a complete understanding of the downstream effects elicited by TFs and diverse chromatin-associated factors. To address this gap, we have established a platform to systematically identify and characterize the transcriptional regulatory potential of human proteins in an unbiased manner. By screening over 13,000 proteins in a pooled format, we identified several hundred potent activators, many of which were previously poorly annotated. We also systematically uncovered transactivation domains among the hits, including some that do not adhere to the canonical “acidic blob” model of activation domains. Furthermore, we combined interaction proteomics and chemical inhibitors to delineate the co-factor specificity of both novel and known transcriptional activators, highlighting how even highly related TFs with virtually identical DNA-binding specificities can activate transcription through distinct co-factor complexes.

## RESULTS

### Platform for identifying transcriptional activators in human cells

To identify transcriptional regulators in a systematic manner, we used a chemically-induced dimerization (CID) system, where catalytically inactive Cas9 (dCas9) is tagged in its N terminus with the protein phosphatase ABI1 and the potential transcriptional activator is fused to the abscisic acid receptor PYL1 (**Figure 1A**)(Gao et al., 2016; Liang et al., 2011). Treatment of cells with abscisic acid (ABA) induces an interaction between ABI1 and PYL1, thereby recruiting the potential transcriptional activator to a specific genomic locus defined by the guide RNA (gRNA). As a reporter, we used a HEK293T cell line that contains a stably integrated construct with a 7xTetO array and a basal CMV promoter driving the expression of EGFP (Gao et al., 2016). We generated by lentiviral infection a monoclonal cell line that expresses ABI1-dCas9 and a single gRNA targeting TetO. Transfection of this cell line with known transcriptional activators VPR, VP64, and p300 fused to PYL1 led to robust induction of GFP upon ABA treatment (**Figure 1B**), consistent with a previous report (Gao et al., 2016). Recruitment of luciferase or RFP to the promoter did not induce GFP expression, demonstrating that ABA alone does not activate the reporter (**Figure 1B**).

**Figure 1.**
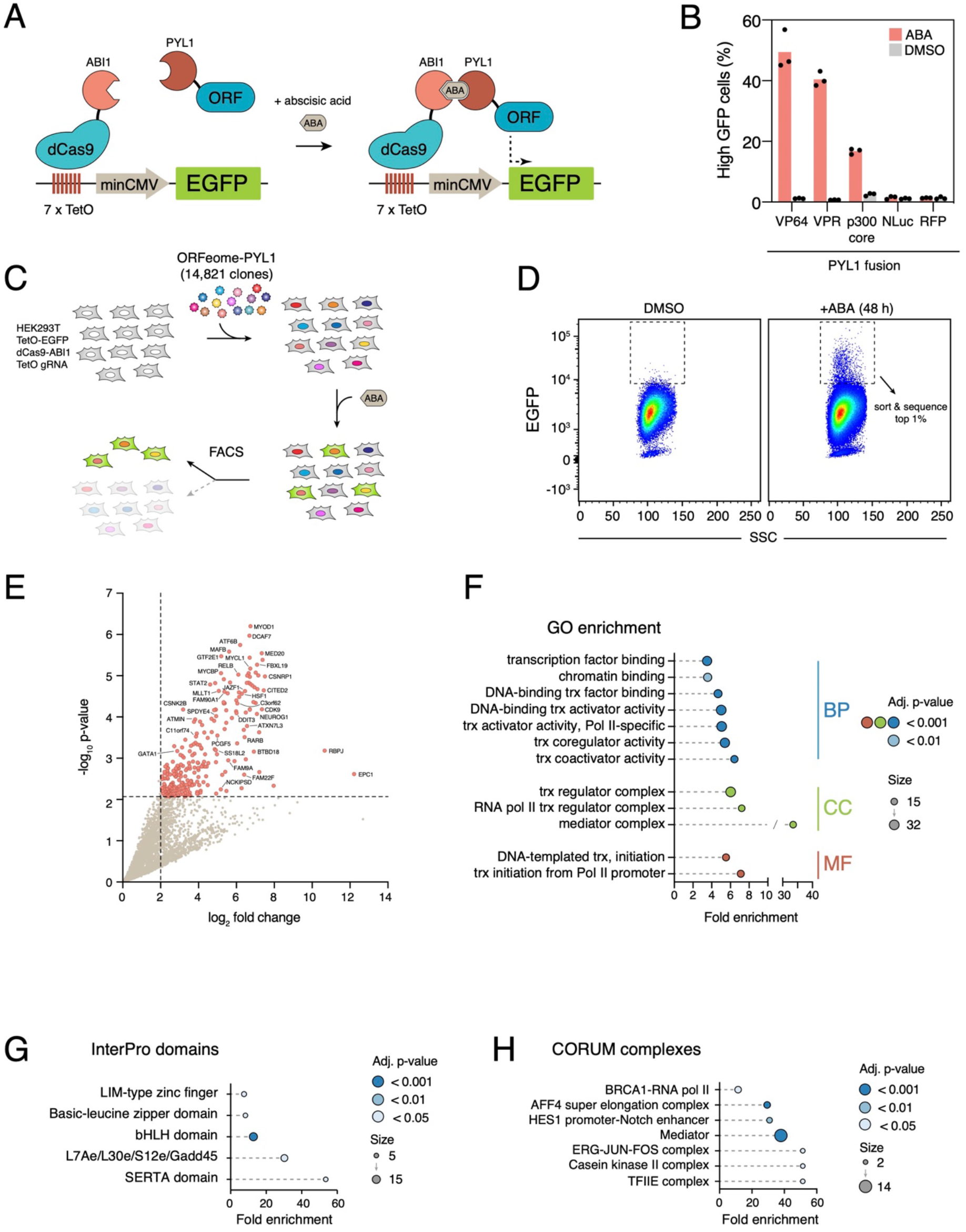
Pooled ORFeome screen for transcriptional activators. **A,** Chemically-induced dimerization system to characterize transcriptional activators in human cells. **B,** Indicated constructs were transfected into HEK293T reporter cells and treated with abscisic acid or DMSO for 48 hours. **C,** Outline of the pooled ORFeome screen for transcriptional activators. **D,** Enrichment of high GFP cells in the pooled ORFeome screen after 48 hour ABA treatment. **E,** Enrichment of ORFs in the high GFP pool compared to the unsorted ORFeome. **F,** Enrichment of Gene Ontology categories among positive screen hits. **G,** Enrichment of InterPro domains among positive screen hits. **H,** Enrichment of CORUM complexes among screen hits.

To scale up from individual clones, we generated a pooled library of human ORFeome 8.1 and ORFeome collaboration clones (ORFeome Collaboration, 2016; Yang et al., 2011) in a lentiviral vector containing a C-terminal PYL1. Together, these open reading frame (ORF) libraries contain 14,821 clones corresponding to 13,571 unique genes. We infected the reporter cell line with the pooled library at low multiplicity of infection, thus ensuring that most cells were infected with only one lentivirus (**Figure 1C**). The pooled library covered 96% and 93% of the ORFeome before and after infection, respectively, demonstrating wide coverage of the library despite significant diversity in insert size (**Figure S1A** and **S1B**). After treating the infected cells with ABA, we noticed a clear induction of GFP in a subset of cells (**Figure 1D**). Reporter induction was dependent on treatment time, ABA concentration, and presence of the ORFeome (**Figures S1C-S1E**). Furthermore, withdrawal of ABA led to rapid loss of high GFP population (**Figure S1C**), demonstrating that continuous promoter occupation is required for activation. To identify transcriptional activators, we sorted top 1% of GFP positive cells of ABA treated cells in duplicate and identified ORFs in the high GFP population by sequencing (**Figure 1D**).

### ORFeome-wide screen identifies known and novel transcriptional activators

We identified 248 putative transcriptional activators, using a cut-off of 5% false discovery rate and at least 4-fold change in read counts between top 1% GFP positive cells and unsorted cells (**Figure 1E** and **Table S1**). To characterize the biological relevance of the hits, we first asked if they represented specific classes of genes. Indeed, gene ontology (GO) analysis revealed significant enrichment for multiple functional categories related to transcriptional activation (**Figure 1F**). The hits were also highly enriched in protein domains found in many transcriptional regulators (**Figure 1G**), subunits of chromatin-associated protein complexes (**Figure 1H**), and in interactors of central hubs of transcription, such as RNA polymerase II and histone acetyltransferases CBP and p300 (**Figure S1F**). Moreover, the hits were significantly overlapping with human proteins that function in yeast two-hybrid assays as autoactivators (11% in the proteome vs. 29% among the hits; p<0.0001, Fisher’s exact test; **Figure S1G**), which are proteins that activate reporter gene expression in yeast when ectopically recruited to the promoter (Luck et al., 2020). We also validated our screen results by assaying a collection of 90 hits and non-hits by individually transfecting them into the same reporter cell line. The results were highly consistent with the screen: almost all hits reproduced when tested individually, and conversely, most non-hits did not activate the reporter (**Figure S1H**), suggesting low false positive and false negative rates in this setting. Together, these results indicate that the screen identified functionally relevant transcriptional activators in a reproducible manner.

Individual screen hits included well-characterized factors regulating distinct stages of transcription (**Figure 1E** and **Table S1**). For example, among sequence-specific transcription factor hits were known activators RELB and MYCL (Barrett et al., 1992; Ryseck et al., 1992) in addition to several master TFs regulating stress response, such as HSF1, ATF6, and DDIT3/CHOP (Vihervaara et al., 2018). Co-activators that do not bind DNA themselves but associate with TFs included STAT2, CITED1, and SERTAD1 (Bousoik and Montazeri Aliabadi, 2018; Hsu et al., 2001; Yahata et al., 2001). Both subunits of TFIIE (GTF2E1 and GTF2E2), a general transcription factor regulating the assembly of the pre-initiation complex (Cramer, 2019), were also prominent hits, as were proteins promoting RNA polymerase II release (the P-TEFb complex subunit CDK9) and transcriptional elongation (e.g., ELL3 and MLLT1)(Chen et al., 2018; Cramer, 2019). Other prominent activators included subunits of chromatin-modifying complexes such as SAGA (ATXN7L3, TADA3, SGF29) and NuA4 (EPC1, MGRBP), and GADD45 family proteins that mediate DNA demethylation (GADD45A, GADD45B, GADD45G)(Barreto et al., 2007). Finally, we identified 13 of the 15 Mediator subunits that were present in the library. These highlighted examples illustrate that the screen uncovered factors promoting transcription by multiple different mechanisms and at distinct stages of the transcription cycle.

In addition to known transcriptional regulators, our screen identified a large collection of proteins that have not been previously linked to transcription (e.g. C11orf74/IFTAP, NCKIPSD, DCAF7, HFM1) or are completely uncharacterized (e.g. C3orf62, FAM90A1, SPDYE4, SS18L2, FAM9A, C21orf58)(**Figure 1H**). These results suggest that the human genome encodes many previously unknown transcriptional regulators. Detailed characterization of several of these factors is described below.

### Distinct transcriptional activities within TF families

Transcription factors comprise multiple families, characterized by their distinct DNA-binding and auxiliary domains (Lambert et al., 2018). TFs that belong to the same family often have highly similar or even identical sequence specificities (Jolma et al., 2013; Lambert et al., 2018; Weirauch et al., 2014). Nevertheless, even highly related TFs can have distinct effects on transcription and chromatin due to their unique auxiliary domains. In line with this, we noticed that only some members of transcription factor families were hits in the pooled screen. To rule out sensitivity of the pooled approach as the cause, we individually assayed members of Forkhead-box (FOX), SRY-related HMG-box (SOX), E-twenty-six (ETS), atonal-related basic helix-loop-helix (bHLH), Twist/Hand, Krüppel-like factor (KLF), and Homeobox protein (HOX) families (**Figure 2A** and **S2A**). We also included two protein families linked to transcription and chromatin (Polycomb group RING finger (PCGF) and casein kinase family), one family (Spy1/RINGO) not previously linked to transcription (Gastwirt et al., 2007), and 24 Mediator subunits, which are not evolutionarily related but are integral components of the same, conserved complex.

**Figure 2.**
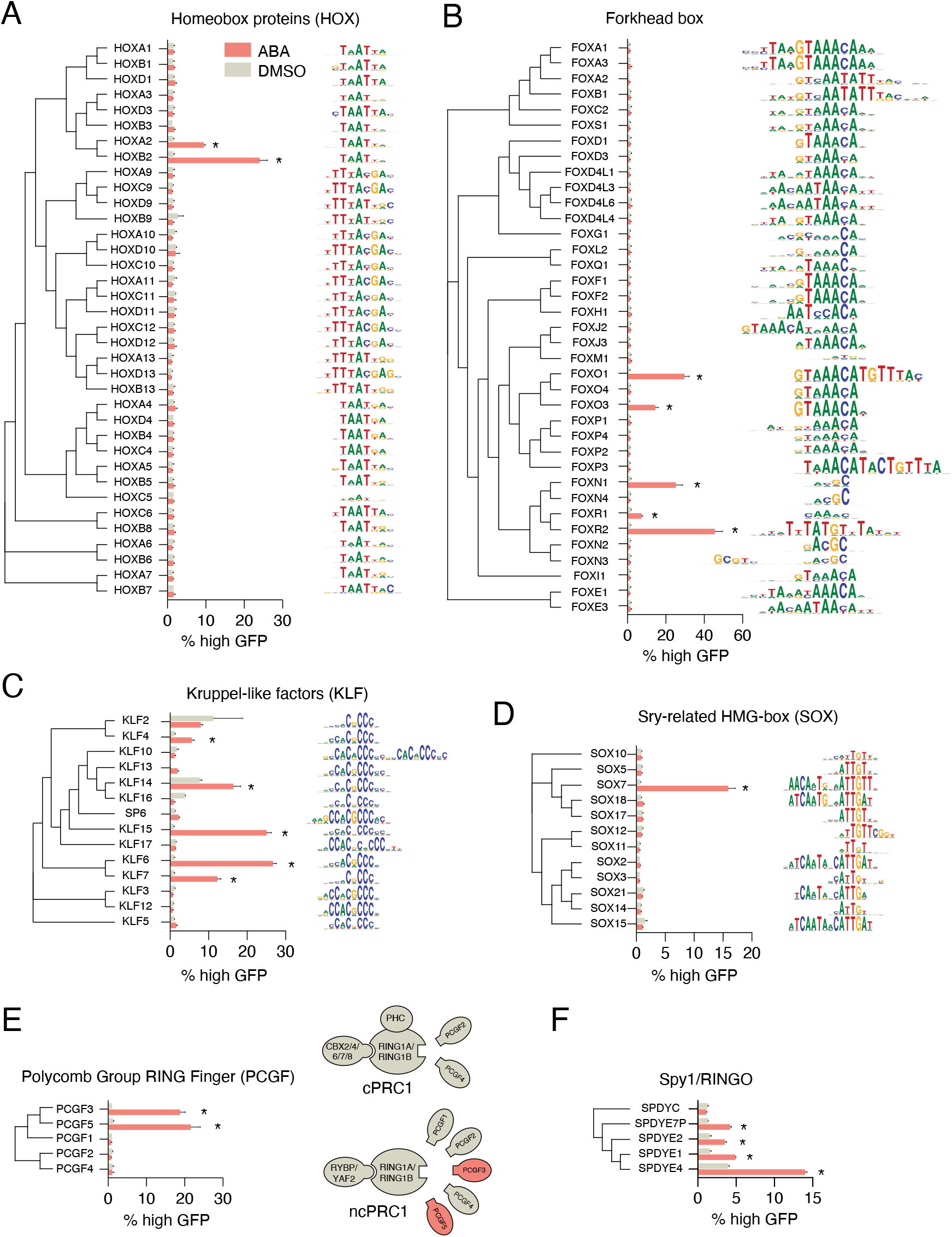
Transcriptional activity of transcription factor families. Transcriptional regulators were individually tested for activation of the reporter in an arrayed manner. DNA-binding specificity (shown as sequence logos) is from CisBP (Weirauch et al., 2014). Asterisks indicate statistically significant activators (FDR < 0.05). **A,** Homeobox family proteins **B,** Forkhead box proteins, **C,** Kruppel-like factors, **D,** SRY-related HMG-box (SOX) proteins, **E,** Polycomb-group RING Finger (PCGF) proteins. Composition of canonical (cPRC1) and non-canonical (ncPRC1) complexes is shown on the right. **F,** Spy1/RINGO family proteins. Statistical significance was calculated with unpaired two-tailed t-test assuming equal variance, and corrected for multiple hypotheses with False Discovery Rate (FDR) approach of Benjamini, Krieger and Yekutieli (Benjamini et al., 2006).

Consistent with the pooled screen, activation profiles for many highly homologous transcription factors were markedly different even when tested one at a time. For example, only five of the 37 Forkhead TFs, one of the 14 SOXs, five of 14 KLFs, and two of the 36 HOX proteins activated reporter expression (**Figure 2A-2D** and **S2**). To exclude the trivial explanation that the difference in activity was due to expression, we assayed several TFs as RFP-PYL1 fusions to account for fusion protein expression levels. The results were highly consistent with PYL1-only fusions and did not correlate with RFP levels (**Figure S3**), indicating that the differences in transcriptional activity reflected the intrinsic potential of the TFs.

Notably, activating TFs in our screen were significantly enriched for factors that can induce differentiation of mouse embryonic stem cells and human induced pluripotent cells (iPSCs) when ectopically expressed (**Figure S2G**)(Ng et al., 2021; Theodorou et al., 2009), suggesting that activation differences in our assay are related to differences in biological function. For example, of the related bHLH family transcription factors, NEUROG1, NEUROG2, NEUROD1, and NEUROD2 can induce neuronal differentiation of iPSCs whereas NEUROD6 cannot (Goparaju et al., 2017). This pattern is concordant with their ability to activate the reporter gene (**Figure S2B**). Similarly, only four HOX factors (HOXA1, HOXA2, HOXB1, HOXB2) can activate the b1-ARE autoregulatory element located in the *Hoxb1* locus (Di Rocco et al., 1997), and our assay characterized three of these as activators (HOXA2 and HOXB2 in the pooled screen and the arrayed assay, HOXA1 in the screen) (**Figure 2A**). Moreover, recent work revealed a striking collinearity between the repressive potential, expression pattern, and genomic location of HOX genes (Tycko et al., 2020), with HOX transcription factors in the 5’ end of homeobox clusters being repressive. Consistent with this, the HOX family activators in our assay are encoded by the most or the next-to-last 3’ genes in their HOX gene clusters.

A particularly interesting case was that of PCGF family proteins, which are mutually exclusive components of canonical and non-canonical Polycomb Repressive Complexes 1 (PRC1) (Gahan et al., 2020; Gao et al., 2012)(**Figure 2E**). Although generally thought to act in chromatin compaction and gene silencing in the context of PRC1, we identified PCGF3 as an activator in the original screen. PCGF3 robustly activated the reporter when tested individually, as did PCGF5 (which was not present in the original pooled screen) **(Figure 2E)**. In contrast, three other PCGF family members (PCGF1, PCGF2, and PCGF4/BMI1) were neither screen hits nor activated the reporter when individually tested (**Figure 2E**). PCGF5 has been previously shown to regulate transcriptional activation (Gao et al., 2014), but no such role has been described for PCGF3. Interestingly, in the PCGF family phylogeny, PCGF3 and PCGF5 form a distinct group that arose early during animal evolution (Gahan et al., 2020), suggesting that their transcriptional activation function is of ancient origin.

Some studies have indicated that subunits of the Mediator complex can have distinct regulatory functions, with some subunits promoting transcriptional activation and others promoting repression (Conaway and Conaway, 2011; Stampfel et al., 2015). In our assay, 20 of the 24 assayed Mediator subunits robustly activated the reporter, with no difference between Mediator submodules (**Figure S2F**). For example, MED29 has been suggested to have transcriptional repressor activity (Wang et al., 2004), but it was both a hit in the pooled screen and validated as an activator when tested individually (**Figure S2F**). Thus, at least in the context of our reporter system, almost all Mediator subunits promote transcriptional activation.

The primary activation screen identified two proteins (SPDYE4 and SPDYE7P) that belong to the Spy1/RINGO (Rapid INducer of G2/M progression in Oocytes) family of cell cycle regulators. Spy1/RINGO proteins bind to and activate Cdk1 and Cdk2 in a cyclin-independent manner, thereby promoting cell cycle progression (Gonzalez and Nebreda, 2020). However, they have not been previously implicated in transcriptional regulation. As a result of recent expansion (Chauhan et al., 2012), the human genome contains at least 19 Spy1/RINGO family genes and multiple pseudogenes. We individually tested five Spy1/RINGO proteins for transcriptional activation, and four of them robustly activated the reporter (**Figure 2F**). These results suggest that Spy1/RINGO proteins may promote cell cycle progression in part by functioning as transcriptional activators.

### TAD-seq reveals novel human transactivation domains

Transcription factors generally activate transcription through transactivation domains (TADs) that interact with co-activators, such as the Mediator, CBP/p300 acetyltransferases, or TFIID. Most TADs are short, unstructured sequences rich in acidic and hydrophobic residues (Sigler, 1988). Despite intensive efforts, no clear consensus motifs have emerged, making computational prediction of TADs challenging (Erijman et al., 2020; Ravarani et al., 2018; Staller et al., 2021). Several groups have recently implemented pooled recruitment approaches to identifying TADs from known transcriptional regulators or from random sequences (Arnold et al., 2018; Erijman et al., 2020; Ravarani et al., 2018; Sanborn et al., 2020). We modified our screening platform to identify the region(s) responsible for transcriptional activation among our hits. Our approach was similar to the previously published TAD-seq method (Arnold et al., 2018) except that we used synthesized fragments instead of randomly fragmented DNA.

We generated a fragment library of 75 activators identified in our screen, using 60 amino-acid tiles every 20 aa such that every amino acid was represented by three different fragments (**Figure 3A**). We fused the pooled fragment library to PYL1 and recruited it to the same GFP reporter used in the original ORFeome screen (**Figure 3A**). Again, we observed ABA-dependent GFP expression in a subset of reporter cells infected with the fragment library (**Figure S4A**). We sorted the GFP positive population into two independent replicates, and quantified fragments in each population by sequencing. However, in contrast to the ORFeome screen, we sorted both high GFP cells (top 1%) and medium GFP cells (top 2-5%) to potentially identify transactivators of different strength.

**Figure 3.**
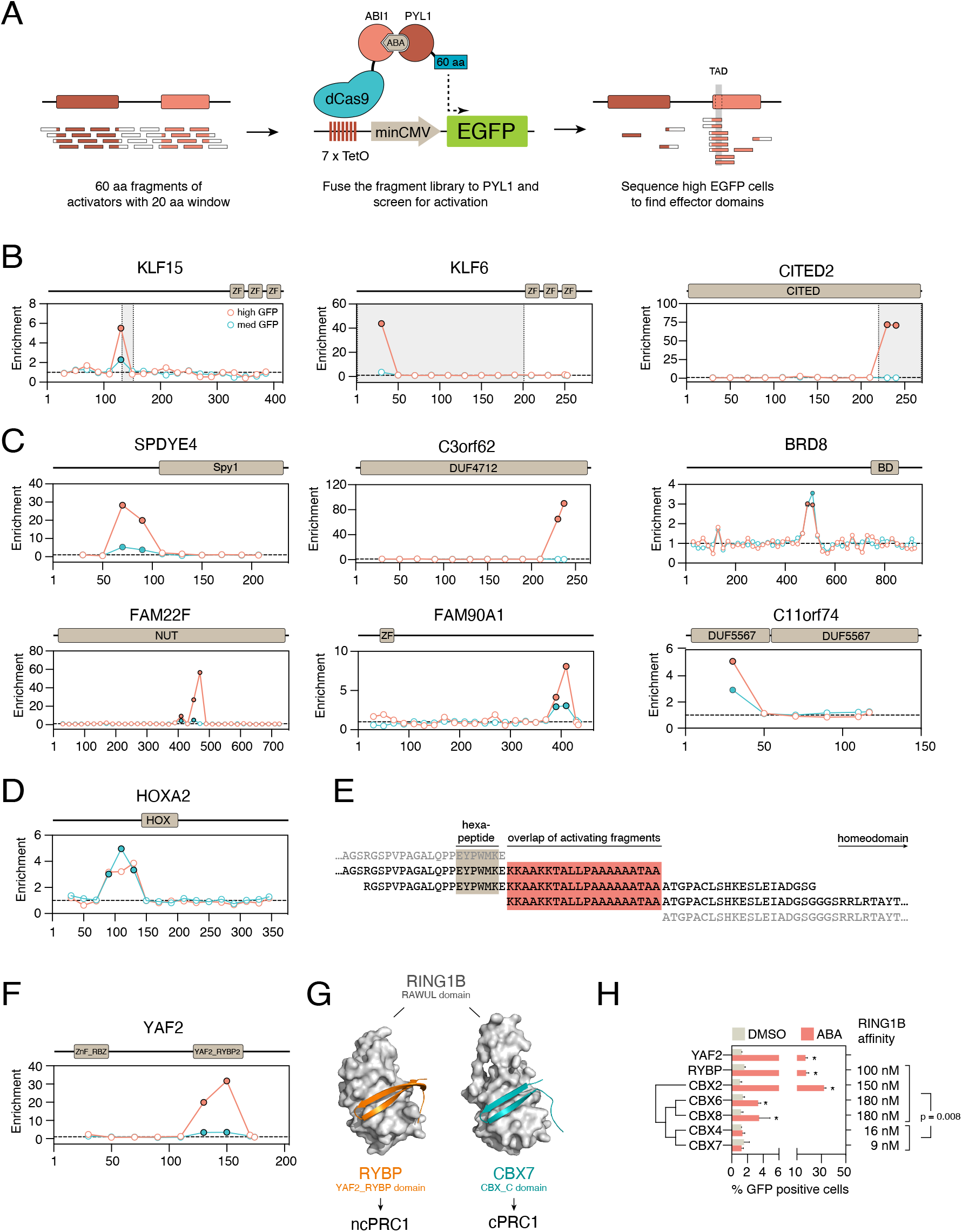
Systematic discovery of transactivation domains in human proteins with TAD-seq. **A,** Outline of the TAD-seq pooled assay. **B,** Examples of known transactivation domains. Domain organization is shown on top. TAD-seq plot shows the fold enrichment of RNAseq reads in the high GFP population (red) or the medium GFP population (blue). Each circle shows the mid-point (30th amino acid) of the 60-aa tile. Filled circles indicate statistically significant hits. Grey boxes indicate previously described transactivation domains. **C,** Examples of novel transactivation domains. Labeling is as in panel B. **D,** Location of the transactivation domain of HOXA2. **E,** Sequences of the activating fragments identified in HOXA2. Activating fragments are in bold. Red box indicates the region common to all three fragments that were enriched in the medium GFP population. Light brown box indicates the location of the antennapedia-like hexapeptide sequence. **F,** Location of the transactivation domain in YAF2. **G,** Crystal structures of RING1B RAWUL domain bound to the YAF2_RYBP domain of RYBP (orange; PDB 3IXS) and the CBX_C domain of CBX7 (cyan; PDB 3GS2). **H,** YAF2_RYBP domains and CBX_C domains from indicated proteins were individually tested for transcriptional activity. Asterisks indicate statistically significant activators (FDR < 0.05). Statistical significance was calculated with unpaired two-tailed t-test assuming equal variance, and corrected for multiple hypotheses with False Discovery Rate (FDR) approach of Benjamini, Krieger and Yekutieli (Benjamini et al., 2006). Right, *in vitro* affinity of the domains towards RING1B (Wang et al., 2008, 2010). Statistical significance was calculated with an unpaired two-tailed t-test.

The pooled approach revealed 70 activating fragments in 39 different proteins (**Table S1**). As expected, these fragments were enriched in acidic and hydrophobic amino acids, and depleted of positively charged (basic) amino acids (**Figure S4B**), indicating that many represent “canonical” TADs consistent with the “acid blob” idea (Sigler, 1988). Indeed, activating fragments contained more predicted TADs based on two different algorithms (Erijman et al., 2020; Piskacek et al., 2007)( **Figure S4C**) and the predicted TADs were longer in activating fragments than in inactive fragments (**Figure S4D**). However, many active fragments were not predicted by any algorithm, highlighting the need for experimental approaches.

Of the 70 active fragments, the majority (44 = 63%) was identified in only the high GFP or the medium GFP population, suggesting that the fragments have distinct activation potential (**Figure S4E**). For example, VP64 was enriched in the high GFP population but depleted in the medium GFP population when compared to the unsorted fragment library (**Figure S4E**). This is consistent with the unusual potency of VP16 in transcriptional activation (VP64 consists of four VP16 peptides in tandem)(Sadowski et al., 1988). We individually tested 10 fragments identified in the screen, and 9 of them robustly activated the reporter (**Figure S4F**), suggesting low false positive rate.

The fragment screen recovered several known TADs. For example, we identified known TADs in KLF6, KLF7, KLF15, ATF6, and CITED2 (**Figure 3B** and **S5A**). Additionally, since the overlap between active fragments can pinpoint the minimum sequence required for activity, we were able to narrow regions required for activity of previously identified TADs, such as those for KLF6, KLF7, and SERTAD2 (**Figure 3B** and **S5A**). We also identified many previously unknown TADs in both canonical transcriptional regulators and novel proteins we discovered in the ORFeome-wide screen (**Figure 3C** and **S5B**). TADs were uncovered in previously uncharacterized proteins SPDYE4, C3orf62, FAM22F, and FAM90A1 (**Figure 3C**). Interestingly, the novel TAD in SPDYE4 was adjacent to its Spy1 domain that interacts with and activates Cdk2, indicating that the potential transcriptional activity and Cdk-regulating activity are mediated by distinct domains of Speedy family proteins (**Figure 3C** and **S6A**)(McGrath et al., 2017). Moreover, consistent with the transcriptional activation profiles of Spy1/RINGO family proteins, the TAD region is conserved in all family members except SPDYC, the only Spy1/RINGO family protein that was inactive in the recruitment assay (**Figure 2F** and **S6A**).

Interestingly, some of the uncovered TADs did not have characteristics of typical transactivation domains. For example, the three overlapping fragments in HOXA2 that activated transcription spanned a polyalanine stretch between the homeobox DNA-binding domain and the antennapedia-like hexapeptide motif (**Figure 3D** and **3E**), which interacts with the PBX1 co-activator (Piper et al., 1999). However, a fragment lacking the hexapeptide motif still activated transcription, suggesting that the activity is not regulated via PBX1 (**Figure 3E**). Polyalanine stretches are functionally important in Ultrabithorax (Ubx) proteins in *Drosophila*, and recently they were suggested to have a role in driving phase separation of many TFs such as HOXD13 (Basu et al., 2020). Our result suggests that polyalanine stretches may have additional roles in transcriptional activation.

Another non-canonical activating region was from YAF2, which is a component of the Polycomb Repressive Complex 1 (PRC1) (Gao et al., 2012) and a prominent hit in the original ORFeome screen (**Figure 3F** and **Table S1**). This region contained the YAF2_RYBP domain, which folds into an antiparallel beta sheet that binds RING1B, a core PRC1 subunit (Wang et al., 2010) (**Figure 3G**). The YAF2_RYBP domain of YAF2 and its close homolog RYBP shares sequence and structural homology with the CBX-C domain present in the CBX family Polycomb proteins (Wang et al., 2010)(**Figure S6B**). CBX-C domain containing proteins and YAF2/RYBP interact in a mutually exclusive manner with RING1A and RING1B to form canonical (cPRC1) and non-canonical (ncPRC1) Polycomb complexes (**Figure 3G**). Intriguingly, the YAF2_RYBP domain was recently shown to be sufficient to induce silencing when recruited to an active promoter (Tycko et al., 2020), indicating that the domain can function both as a repressor and an activator even in relatively simple reporter systems.

To test if all CBX-C and YAF2_RYBP domains can promote transcriptional activation, we cloned the YAF2_RYBP domains of YAF2 and RYBP and the CBX-C domains of CBX2, CBX4, CBX6, CBX7, and CBX8 and assayed their activity with the reporter system. The YAF2_RYBP motif from both YAF2 and RYBP robustly activated the reporter, consistent with the TADseq results for YAF2 (**Figure 3H**). Some CBX-C domains were also activators: CBX-C from CBX2 was the most potent activator, whereas those from CBX6 and CBX8 activated the reporter weakly (**Figure 3H**). In contrast, CBX4 and CBX7 CBX-C domains had no effect on the reporter. This pattern reflected the evolutionary ancestry of the CBX-C domain proteins (**Figure 3H**). Interestingly, the CBX-C domains of CBX4 and CBX7 bind RING1B with ∼10-fold higher affinity than the same domains from CBX proteins that activate transcription, or the YAF2_RYBP domain of RYBP (p = 0.008, two-tailed t-test)(**Figure 3H**)(Wang et al., 2008, 2010). Thus, the differences in the transcriptional activity of CBX-C and YAF2_RYBP domains might be explained by their binding affinity for RING1B or by differential binding to other factors. In any case, our results suggest that the differences in CBX protein function (Morey et al., 2012; Vincenz and Kerppola, 2008) could be at least partially explained by the intrinsic differences in their CBX-C domains. More broadly, these results reveal another layer of complexity in the assembly and function of noncanonical and canonical PRC2 complexes.

### Novel transcriptional activators interact with known co-factors

The ORFeome-wide screen revealed several potent transactivators that were either poorly or completely uncharacterized. To understand how these factors regulate transcription, we established stable, tetracycline-inducible HEK293 cell lines expressing nine poorly characterized screen hits (C3orf62, C11orf74/IFTAP, NCKIPSD, DCAF7, SS18L2, SPDYE4, FAM90A1, FAM22F/NUTM2F, JAZF1), five known transcriptional regulator hits (SOX7, KLF6, KLF15, CTBP1, HOXA2, and HOXB2), two synthetic transactivators (VP64 and VPR), and negative controls (EGFP and Nanoluc) fused to biotin ligase BirA from *Aquifex aeolicus* and FLAG epitope tag. We characterized their protein interactomes with affinity-purification coupled to mass spectrometry (AP-MS) and proximity partners with proximity-dependent biotinylation (BioID2)(Kim et al., 2016). AP-MS is an ideal method for characterizing stable protein complexes, whereas BioID excels in identifying interactions that are weaker or involve poorly soluble proteins, such as those tightly bound to chromatin (Lambert et al., 2015).

The interactomes of transcriptional activators revealed two patterns. First, they indicated that the transactivation potential of the novel hits likely reflects their endogenous function rather than being an artefact of the tethering assay. Second, the interactomes uncovered a striking preference of the activators for specific co-activator complexes, converging on five distinct co-factors (CBP/p300, BAF, NuA4, Mediator, and TFIID).

Supporting a native role for the novel activators in transcriptional regulation, eight of the nine poorly characterized hits associated with known transcriptional co-factors in AP-MS, BioID, or both (**Figure 4A**, **Figure S7** and **Table S1**). C3orf62, DCAF7, and FAM22F/NUTM2F interacted with p300 and/or CBP, which are known transcriptional co-activators. JAZF1, in contrast, associated with multiple subunits the NuA4 histone acetyltransferase complex in BioID, suggesting that it is a novel subunit of this highly conserved complex. SS18L2, in turn, interacted with the BAF chromatin remodeling complex, including core BAF members SMARCA2 and SMARCA4 as well as subunits specific to the canonical BAF (cBAF) and non-canonical BAF (ncBAF) (**Figure 4A** and **S7**)(Centore et al., 2020). SPDYE4 and FAM90A1 interacted with the BET family bromodomain proteins BRD2, BRD3, and BRD4, which regulate transcriptional elongation (Fujisawa and Filippakopoulos, 2017). NCKIPSD, also known as SPIN90, interacted with the survival of motor neurons (SMN) complex, which regulates the assembly of ribonucleoprotein complexes but has also been linked to transcriptional activation (Pellizzoni et al., 2001; Singh et al., 2017; Strasswimmer et al., 1999) (**Figure S7**). Interestingly, NCKIPSD also interacted with DCAF7 (**Figure S7B**). Both DCAF7 and NCKIPSD have been implicated in the regulation of actin dynamics in the cytoplasm (Cao et al., 2020; Morita et al., 2006), suggesting that these proteins may have distinct roles in the cytoplasm and in the nucleus.

**Figure 4.**
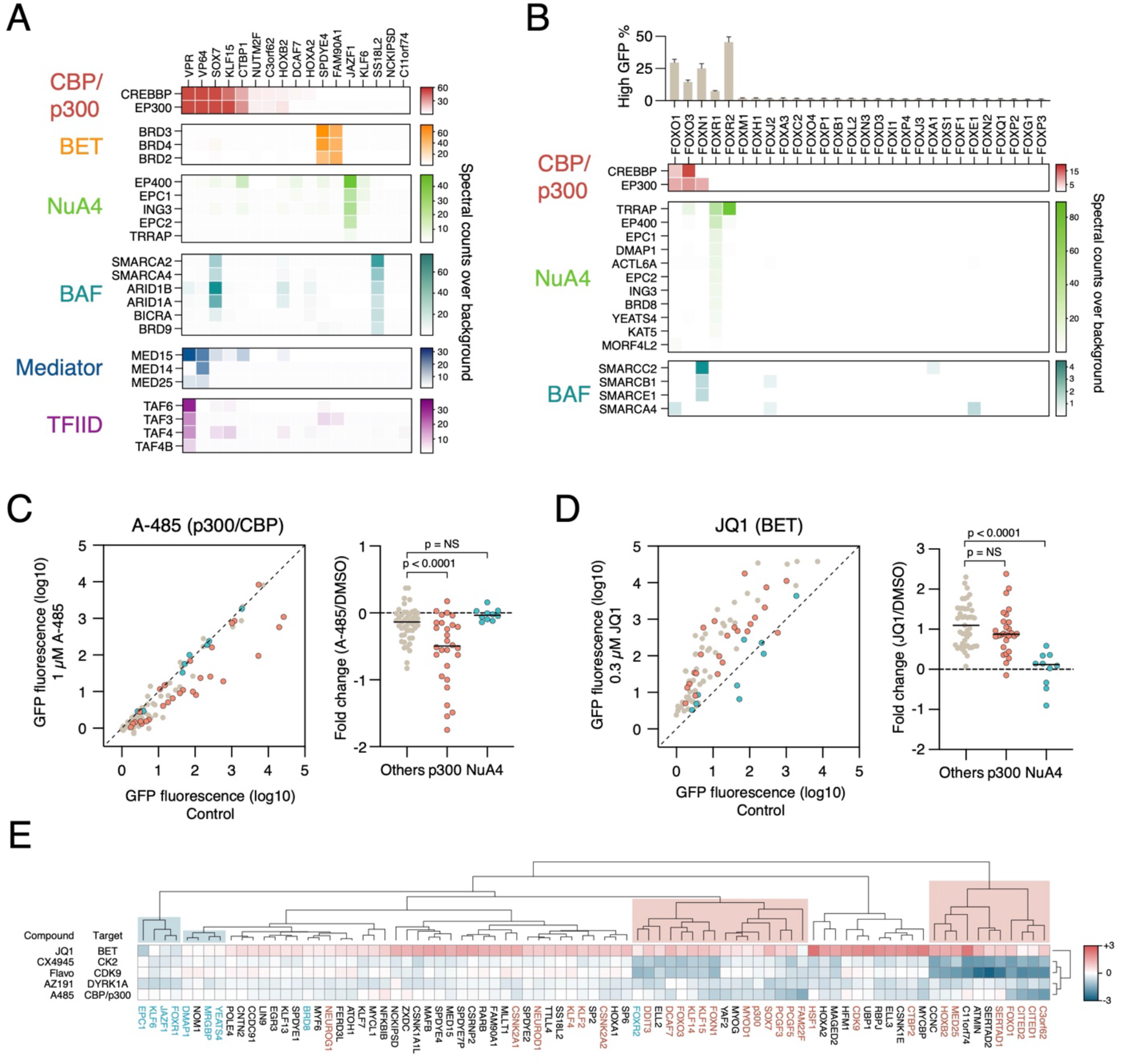
Co-factor specificity of transcriptional activators. **A,** Proximity partners of indicated transcriptional regulators were identified with BioID2. Enrichment of selected co-factor complexes is shown as a heat map. Average spectral counts (n+1) were normalized to background spectral counts (n+1) of EGFP and Nanoluc baits. **B,** Interaction patterns of activating Forkhead transcription factors based on the AP-MS study of (Li et al., 2015). Spectral counts were normalized as in panel A. **C,** Left, effect of p300/CBP inhibition by A-485 on the activity of 83 transcriptional regulators. Known p300 interactors are shown as red circles, known NuA4 interactors as blue circles. Right, p300 interactors are significantly more affected than other transcriptional regulators by A-485 treatment. Statistical significance was calculated with one-way ANOVA using Dunnett’s multiple testing correction. **D,** Left, effect of BET bromodomain protein inhibition by JQ1 on the activity of 83 transcriptional regulators. Known p300 interactors are shown as red circles, known NuA4 interactors as blue circles. Right, p300 interactors are significantly more affected than other transcriptional regulators by A-485 treatment. Statistical significance was calculated with one-way ANOVA using Dunnett’s multiple testing correction. **E,** Clustering of transcriptional activators based on their sensitivity on diverse inhibitors. Clustering was performed with Euclidian distance with average linkage. Clusters enriched in known CBP/p300 interactors (red) and NuA4 interactors (blue) are highlighted.

Known transcriptional regulators also associated with co-activator complexes (**Figures 4A** and **S7** and **Table S1**). KLF6 bait identified the NuA4 subunit EPC1 as a proximity interactor, whereas HOXB2, KLF15, CTBP1, and SOX7 baits had CBP and p300 as proximity partners. In addition, SOX7 also associated with BAF subunits. In contrast to all native activators except for SOX7, the powerful synthetic activators VP64 and VPR identified multiple different co-activators as proximity partners. VP64, which consists of four tandem copies of the viral VP16 transactivation motif, associated with CBP/p300 and mediator subunits MED14 and MED15, consistent with previous reports (Kundu et al., 2000; Yang et al., 2004). VPR, which is a fusion of VP64, human p65/RELA transactivation domain, and Epstein-Barr virus R transactivator (Chavez et al., 2015), associated with even more co-factors, including CBP/p300 and multiple subunits of the Mediator and TFIID (**Figures 4A** and **S7A** and **Table S1**). Such association with multiple co-factors likely explains the exceptional transactivation potency of VPR when fused to dCas9 (Chavez et al., 2015, 2016).

Our results suggest that activating transcription factors and other transcriptional regulators have a strong intrinsic preference for specific co-factors. To further investigate this, we analyzed a previously published AP-MS interaction dataset of Forkhead family TFs (Li et al., 2015) and compared it with the transcriptional activation results in our assay. Notably, only those Forkhead TFs that activated transcription when recruited to the reporter interacted with co-activators in AP-MS (**Figure 4B**). In addition, similar to our mass spectrometry results, activating Forkhead TFs had distinct co-factor preferences: FOXO1 and FOXO3 interacted specifically with CBP and p300, FOXN1 interacted with both p300 and subunits of the BAF complex, whereas FOXR1 and FOXR2 preferred the NuA4 complex (**Figure 4B**). These data strongly suggest that related transcription factors, which recognize highly similar sequences and activate transcription nearly to the same extent, can promote transcription through distinct co-activator complexes.

To functionally investigate the connection between transcriptional activators and co-factors, we arrayed a panel of 83 robust activators (**Table S1**). We then tested their activation potential using the reporter assay, but now in the presence of small-molecule inhibitors targeting multiple transcriptional co-regulators. We employed three kinase inhibitors targeting transcriptional kinases (flavopiridol for CDK9, CX-4945 for casein kinase 2, and AZ191 for DYRK1A and DYRK1B), and two compounds inhibiting transcriptional co-factors (A-485 for CBP/p300, and JQ1 for BET family bromodomain proteins).

The three kinase inhibitors affected nearly all activators, although to a different degree. Inhibiting CDK9 with flavopiridol led to nearly complete loss of activity of all activators, consistent with the key role of p-TEFb in promoter clearance (**Figure S8A**). Inhibition of casein kinase 2, which regulates transcriptional elongation (Basnet et al., 2014), also led to a general albeit modest attenuation of transcriptional activation (**Figure S8B**). DYRK1A/DYRK1B inhibition had a more subtle effect than the other two compounds, slightly attenuating the activity of most activators (**Figure S8C**). Interestingly, two activators that were not affected by the DYRK1A/DYRK1B inhibitor AZ191 were DCAF7, which forms a conserved complex with DYRK1A (Breitkreutz et al., 2010; Yu et al., 2019), and NCKIPSD, which interacts with DCAF7 (**Figure S8C**).

In contrast to kinase inhibitors that had broad effects on transcription, inhibiting the acetyltransferase activity of CBP/p300 strongly affected the activity of some but not all transactivators (**Figure 4C**). Importantly, these effects were consistent our AP-MS and BioID results: the activity of four of the six CBP/p300 interactors was significantly decreased, whereas only one of the seven non-interactors was affected by A-485 (**Figure S8D**). For example, A-485 inhibited the activity of KLF15, whereas it had no effect on the related Kruppel-like factor KLF6 (**Figure S8D**). More broadly, the activity of proteins known to interact with CBP/p300 was significantly more inhibited by A-485 than that of non-interacting transactivators (**Figure 4C**). In comparison, interactors of NuA4 complex were not significantly affected by CBP/p300 inhibition (**Figure 4D** and **S8D**).

Similar to CBP/p300 inhibition, BET family inhibition with JQ1 had distinct effects on some transactivators. Interestingly, in most cases JQ1 treatment led to an increase in reporter gene activity (**Figure 4D**). Although JQ1 treatment leads to rapid downregulation of BRD4 target genes in many cases (Lovén et al., 2013; Muhar et al., 2018), it has also been linked to transcriptional activation of reporter genes (Sdelci et al., 2016), which likely explains the effect we observed. Nevertheless, not all transactivators responded similarly to JQ1 treatment. In particular, factors interacting with subunits of NuA4 complex were not affected by JQ1 treatment (e.g., JAZF1 and KLF6; **Figure 4D** and **S8D**). Indeed, NuA4 interactors were significantly less affected by JQ1 treatment that CBP/p300 interactors or other transactivators (**Figure 4D**). Although the mechanism by which NuA4 interactors respond to JQ1 treatment in a unique manner requires further research, these results demonstrate how transcriptional activators promote transcription via different co-activator complexes in the context of a single promoter. Indeed, hierarchical clustering of the activators based on their sensitivity to the five compounds revealed multiple distinct groups (**Figure 4E**). Many paralogous factors (such as CITED1 and CITED2, or PCGF3 and PCGF5) clustered next to each other, indicating that clustering produced functionally relevant groups. Moreover, known CBP/p300 interactors were primarily in two distinct clusters, as were NuA4 interactors (**Figure 4E**). It is likely that other members of these clusters similarly use CBP/p300 or NuA4 as co-activators, such as YAF2 or MYOG for p300, or NOM1 for NuA4.

### SRF-C3orf62 fusion interacts with CBP/p300 and promotes SRF/MRTF transcriptional program

Fusion proteins involving transcriptional regulators are common hallmarks of certain cancers, such as leukemias and sarcomas. We noticed that hits from our ORFeome-wide screen were significantly enriched for genes documented in the COSMIC database (cancer.sanger.ac.uk) as fusion partners in diverse cancers (p = 0.019; hypergeometric distribution test). These included well-characterized fusion partners such as ERG, which is fused to EWSR1 in Ewing sarcoma and to TMPRSS2 in prostate cancer; DDIT3/CHOP, fused to EWSR1 or FUS in myxoid liposarcoma; CRTC1, fused to MAML2 in mucoepidermoid carcinoma; and ENL/MLLT1, fused to MLL in mixed lineage leukemia (**Table S1**). In addition, several hits have been described in literature as fusion partners but not functionally characterized. For example, BTBD18 and NCKIPSD were identified as KMT2A/MLL fusion partners leukemia (Alonso et al., 2010; Sano et al., 2000). Moreover, the fragment of BTBD18 that is fused to MLL contains the transactivation domain we identified by TAD-seq (**Figure S5B**), implicating the TAD in the oncogenic potential of the fusion product. Most MLL fusions involve genes regulating transcriptional elongation, such as the super elongator complex (Winters and Bernt, 2017). Interestingly, BTBD18 has also been shown to promote transcriptional elongation (Zhou et al., 2017).

To gain more insight into the mechanisms by which our activation screen hits might promote tumorigenesis as fusion partners, we focused on two poorly characterized fusions, JAZF1-SUZ12 and SRF-C3orf62. The JAZF1-SUZ12 fusion is a hallmark of low-grade endometrial stromal sarcoma (LG-ESS)(Hrzenjak, 2016), bringing together the Polycomb protein SUZ12 and JAZF1 (**Figure 5A**)(Piunti et al., 2019). SRF-C3orf62 was recently described in a pediatric case of myofibroma/myopericytoma (Antonescu et al., 2017). In this fusion, the DNA-binding domain of Serum Response Factor (SRF) is fused to the C-terminus of C3orf62 (**Figure 5B**). Notably, in both cases the transactivation domain that we identified by TAD-seq (**Figure 3C and S5B**) is retained in the fusion constructs (**Figure 5A** and **5B**)

**Figure 5.**
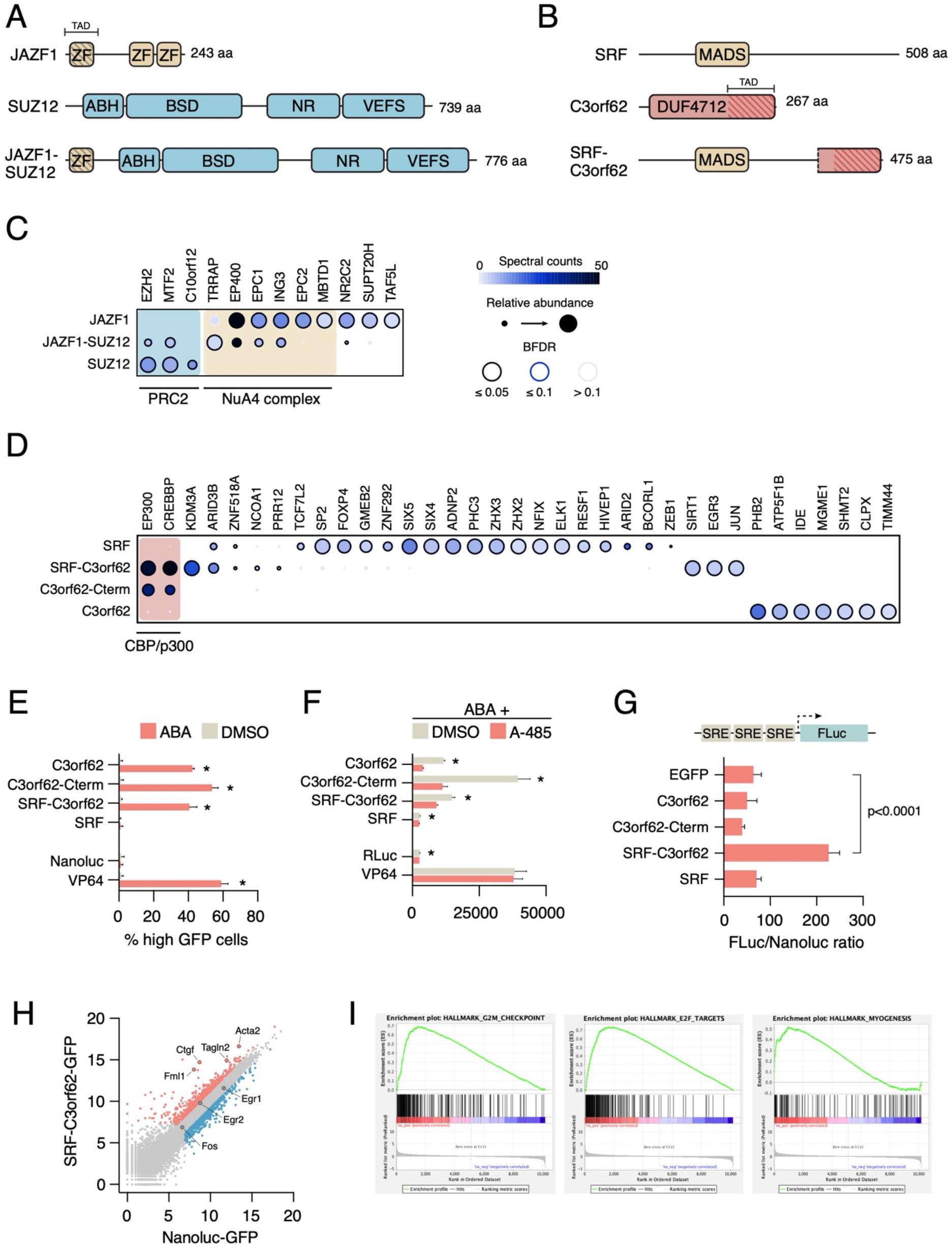
SRF-C3orf62 fusion generates a potent p300-dependent transcriptional activator that promotes expression of SRF/MRTF target genes. **A,** Schematic of the JAZF1-SUZ12 fusion found in low-grade endometrial stromal sarcomas. The transactivation domain identified by TAD-seq is indicated. **B,** Schematic of the SRF-C3orf62 fusion found in myofibroma/myopericytoma. The transactivation domain identified by TAD-seq is indicated. **C,** JAZF1, SUZ12 and JAZF1-SUZ12 proximity interactors were identified with BioID2. JAZF1-SUZ12 interacts with both PRC2 components and NuA4 components, generating a supercomplex. **D,** SRF, C3orf62, C3orf62-Cterm and SRF-C3orf62 proximity interactors were identified with BioID. SRF-C3orf62 fusion robustly interacts with CBP and p300. **E,** SRF-C3orf62 is a potent transcriptional activator. Indicated PYL1 fusions were individually tested for activation of the genomically integrated reporter. **F,** SRF-C3orf62 activates expression of serum response element reporter in the absence of cofactors. Indicated constructs C-terminally tagged with 3xFLAG-V5 were co-transfected into NIH3T3 cells with an SRE-Firefly luciferase reporter and a constitutive Nanoluc reporter. Relative luciferase activities were measured to assess the activity of each construct. **G,** NIH3T3 cells stably expressing doxycycline-inducible SRF-C3orf62-GFP or Nanoluc-GFP were treated with doxycycline for 46 hours, of which the last 22 hours in low-serum conditions (0.5% FCS). Gene expression patterns were analyzed by RNA-seq. Significantly upregulated (red) and downregulated (blue) genes (absolute log2 fold change > 1, FDR < 0.05). Well characterized targets of SRF/MRTF (red, labeled circles) and SRF/TCF (blue, labeled circles) are indicated. **H**, Gene set enrichment analysis of SRF-C3orf62-GFP expressing cells compared to Nanoluc-GFP expressing cells.

We tagged SUZ12, SRF, JAZF1-SUZ12, SRF-C3orf62, and the C-terminal fragment of C3orf62 fused to SRF (C3orf62-Cterm) with BirA-FLAG and analyzed their interactomes with BioID and AP-MS, to complement the data we obtained for JAZF1 and C3orf62. As expected, SUZ12 proximity partners included other components of the PRC2 complex, such as EZH2, MTF2/PCL2 and C10orf12 (Alekseyenko et al., 2014)(**Figure 5C** and **Table S1**). Strikingly, the JAZF1-SUZ12 fusion protein had both PRC2 and NuA4 subunits as proximity partners (**Figure 5C**), indicating that this fusion assembles into a supercomplex of two chromatin-associated complexes that are normally associated with opposing transcriptional activities. Consistent with this, EPC1-PHF1 fusion that is also associated with LG-ESS similarly assembles the PRC2-NuA4 supercomplex, which leads to aberrant expression of Polycomb targets (Sudarshan et al., 2021). Moreover, we have recently shown that both JAZF1-SUZ12 and EPC1-PHF1 are strong transcriptional activators (Sudarshan et al., 2021). Thus, NuA4 integration into the oncogenic supercomplex can override the normally repressive function of PRC2 complexes.

SRF proximity partners included multiple transcription factors such as ELK1, which forms a ternary complex with SRF on serum response elements (SREs)(Buchwalter et al., 2004)(**Figure 5D**). However, it did not associate with any transcriptional co-activators. In contrast, both C3orf62-Cterm construct and the SRF-C3orf62 fusion robustly identified both CBP and p300 as proximity partners (**Figure 5D**). AP-MS similarly identified CBP and p300 as prominent SRF-C3orf62 interactors (**Figure S9A**). Consistent with these results, both C3orf62-Cterm and SRF-C3orf62 strongly activated the GFP reporter when tethered to the reporter (**Figure 5E**). This activity was highly sensitive to CBP/p300 inhibition with A-485 (**Figure 5F**), in line with the interaction patterns of SRF-C3orf62.

SRF functions together with either ternary complex factors (TCFs; e.g. ELK1) or myocardin-related transcription factors (MRTFs; e.g. MAL) to regulate target gene expression (Buchwalter et al., 2004; Olson and Nordheim, 2010). Our results suggested that SRF-C3orf62 can activate SRF target genes without such cofactors. To test this further, we first assayed the activity of different constructs in NIH3T3 fibroblasts using a luciferase-based serum response element reporter (Vartiainen et al., 2007). SRF or C3orf62 alone did not activate the reporter whereas SRF-C3orf62 robustly did so (**Figure 5G**), further confirming that the cancer-associated fusion bypasses the requirement for SRF cofactors in transcriptional activation.

The SRF/TCF pathway, which is regulated by MAP kinase signaling, regulates the expression of immediate-early genes (Gualdrini et al., 2016), whereas the actin-Rho signaling dependent SRF/MRTF pathway targets genes involved in cell motility and adhesion (Miralles et al., 2003). To test if SRF-C3orf62 can regulate target genes of both pathways, we generated stable doxycycline-inducible NIH3T3 cell lines stably expressing SRF, C3orf62, SRF-C3orf62 and Nanoluc fused to GFP. We induced transgene expression with doxycycline for 24 hours and analyzed changes in gene expression by RNA-seq. While expression of GFP-tagged C3orf62 or SRF had very limited or no effects on the transcriptome compared to Nanoluc (**Figure S9B**), expression of SRF-C3orf62-GFP led to upregulation of 564 genes and downregulation of 471 genes (with log2 fold change > 1, FDR < 0.05; **Figure 5H** and **S9B** and **Table S1**). Gene set enrichment analysis (GSEA) and Gene Ontology analysis revealed that the upregulated genes were highly enriched in genes involved in DNA replication, cell cycle progression, and mitosis (**Figure 5I** and **S9C**), whereas downregulated genes were related to extracellular matrix (**Figure S9C**). Upregulated genes were also significantly enriched in targets of the E2F transcription factor, a key regulator of cell proliferation (**Figure 5I**). These signatures are consistent with the oncogenic nature of the SRF-C3orf62 fusion. In addition, many of the most upregulated genes were involved in actin dynamics and myogenesis. For example, one of the most upregulated genes was smooth muscle actin (ACTA2), a known target of SRF/MTRF and a hallmark of SRF fusion positive myofibromas/myopericytomas (Antonescu et al., 2017)(**Figure 5H**). Consistent with this, another significant signature of the upregulated genes was myogenesis (**Figure 5I**). In contrast, many well-characterized immediate-early target genes of SRF/TCF such as FOS, EGR1, or EGR2 were not affected by ectopic SRF-C3orf62 expression (**Figure 5H**). Interestingly, fusion of SRF to VP16 can potently upregulate FOS, suggesting that there are intrinsic differences in the ability of different TADs to activate target gene expression (Schratt et al., 2002). Indeed, there was a significant overlap between genes upregulated by SRF-C3orf62 and previously reported target genes of SRF/MRTF, but not with those of SRF/TCF (**Figure S9D**)(Esnault et al., 2014; Gualdrini et al., 2016). Taken together, these results indicate that SRF-C3orf62 expression leads a proliferative and myogenic gene expression signature and preferential expression of SRF/MRTF target genes over SRF/TCF targets.

## DISCUSSION

Our work presents, to our knowledge, the first systematic screen of transcriptional activators in human cells. We identified several hundred transcriptional activators that were, as expected, enriched in sequence-specific transcription factors and other chromatin-associated proteins. However, it is clear that this does not represent the entire complement of activators in the human genome. For example, the ORFeome collection does not contain many large ORFs (>3000 bp) and thus many chromatin regulators, such as the MLL family, are absent in the collection. In addition, we only used a single core promoter, although it is known that some transcriptional regulators activate transcription only in the context of specific core promoters (Haberle et al., 2019; Stampfel et al., 2015). These limitations could be addressed in further studies using fragment or domain libraries (Arnold et al., 2018; Tycko et al., 2020) and employing different core promoters (Haberle et al., 2019).

Despite these limitations, our screen strongly suggests that only a very limited number of TFs are strong transcriptional activators. This makes sense in the context of the known DNA-binding specificities and chromatin occupancies of human TFs (Jolma et al., 2013; The ENCODE Project Consortium et al., 2020; Yan et al., 2013). Most TFs recognize short (∼6-10 bp) motifs and associate with thousands of sites across the genome. Yet, most binding events are not associated with transcriptional output. Strong transcriptional activation domains coupled to limited sequence specificity of the DNA-binding domain would lead to spurious activation of large swaths of genes, likely impeding cellular fitness. In this context, it is not surprising that many of the strongest activators we identified were not DNA-binding transcription factors but other chromatin-associated proteins.

However, among the strongest activator TFs were master regulators of stress response, such as HSF1, DDIT3, and ATF6. This is in line with their role in heat shock response and unfolded protein response. In normal conditions, their activity is restrained by controlled cytoplasmic localization or chaperone association, but upon stress they must robustly activate their target genes within minutes (Vihervaara et al., 2018). In such cases, strong activation domains are clearly advantageous. Other activating TFs were significantly enriched in factors that can induce differentiation of iPS or ES cells, suggesting that intrinsic activation potential is an important feature of TFs capable of driving cellular reprogramming or differentiation. Indeed, the reprogramming activity of several key transcription factors such as OCT4 or SOX2 can be enhanced by fusing them to more potent activation domains (Hirai et al., 2011; Horb et al., 2003; Narayan et al., 2017; Theodorou et al., 2009; Wang et al., 2011), indicating that transactivation potential can be a limiting factor in cellular reprogramming. Interestingly, MYOD1 and NEUROD1, which can induce transdifferentiation of cells in the absence of any other factors (Davis et al., 1987; Guo et al., 2014), were very potent activators in our screen. We suggest that other strong activator TFs identified in our screen should also be evaluated in cellular reprogramming and differentiation assays.

### Identification of novel transcriptional activators and transactivation domains

Computational prediction of sequence-specific transcription factors is relatively easy due to the conserved nature of DNA-binding domains (Lambert et al., 2018), but prediction of transcriptional activators and repressors is much more challenging due to the degenerate and diverse nature of transcriptional effector domains and motifs. Consistent with this, our functional screen revealed many potent transcriptional activators that have not been previously identified through sequence analysis, candidate-based approaches, or protein/protein interaction studies. These included both completely uncharacterized factors and proteins not previously linked to transcription. For example, we discovered that members of the Speedy/RINGO family of cell cycle regulators are transcriptional activators and contain a potent transactivation domain next to the Spy1 domain that binds to and activates CDKs. Our finding that SPDYE4 also interacts with BET bromodomain proteins such as BRD4 suggests that this it could act as a functional link between CDKs and promoter-proximal pausing of RNA polymerase II during the cell cycle (Core and Adelman, 2019).

Screening fragments of potent activators by TADseq (Arnold et al., 2018) revealed over 70 fragments capable of activating transcription, significantly expanding the catalogue of human TADs. While many of the TADs we discovered contained typical motifs of acidic residues interrupted by hydrophobic amino acids, there were also several surprises. For example, we identified a minimal RING1B-interacting motif in Polycomb group proteins YAF2 and RYBP, as well as in a subset of CBX proteins, that could unexpectedly activate transcription. This finding is particularly surprising, as a recent report identified exactly the same motif as a transcriptional repressor in an analogous screen for repressors (Tycko et al., 2020). Previous work has established that PRC2 complexes have distinct regulatory functions depending on the subunit composition (Gao et al., 2012, 2014). How a simple ∼30aa motif in YAF2 and RYBP can both activate and repress transcription in a context-dependent manner remains to be characterized. Nevertheless, these results suggest an additional layer of regulatory complexity in the assembly of diverse Polycomb complexes.

Our discovery of novel and highly potent human transactivation domains has also therapeutic implications. Components derived from viral proteins, such as the VP16 transactivation domain or the tripartite VPR activator, can elicit immune responses *in vivo* and lead to adverse effects in the clinic. Designing synthetic transcriptional regulators from fully human components would, therefore, be advantageous in therapeutic applications (Israni et al., 2021). Furthermore, it is possible that combining activation domains from multiple different human proteins could generate “superactivators” that are able to robustly upregulate genes even in highly compacted regions of the genome.

### Co-factor specificity, gene regulation, and tumorigenesis

Interactions between transactivation domains and co-factors can be relatively weak and difficult to capture with affinity purification methods. To circumvent this, we employed proximity biotinylation (BioID) with a collection of known and novel activators, as it can detect interactions that would normally be lost during long washing steps. Indeed, BioID revealed a striking specificity of these activators for different co-activators such as CBP/p300, NuA4, and BRD4, suggesting that transcriptional activation can occur through several routes in the context of a single reporter gene. Notably, even closely related activators with almost identical DNA binding motifs could associate with distinct cofactors, as was the case for KLF6 (NuA4) and KLF15 (CBP/p300). Recent work has revealed how transcriptional co-factors have preference for distinct core promoters (Haberle et al., 2019; Stampfel et al., 2015), but which TFs recruit which co-factors to promoters and enhancers is still poorly understood. Our analysis with a limited set of activators suggests that characterizing these TF-cofactor interactions in more detail could help decipher the logic of transcriptional regulation.

Oncogenic rearrangements involve both chromatin-associated proteins and sequence-specific transcription factors. For example, MLL is typically fused to members of the super elongation complex (SEC) in diverse leukemias (Winters and Bernt, 2017), and chromosomal rearrangements fusing NuA4 subunits to PRC2 is a common feature of low-grade endometrial stromal sarcomas (Ali and Rouzbahman, 2015; Sudarshan et al., 2021). These characteristic fusions highlight how activation of oncogenic gene expression programs requires a specific combination of fusion partners. Our results suggest that oncogenic fusions between sequence-specific DNA-binding domains and transactivation domains may similarly be limited to specific combinations. For example, the DNA-binding domain of SRF is fused to NCOA1, NCOA2, or FOXO1 in rhabdomyosarcomas and to RELA or C3orf62 in myofibroma/myopericytoma (Antonescu et al., 2017; Karanian et al., 2020), whereas alveolar rhabdomyosarcoma is characterized by fusions of PAX3 or PAX7 DNA-binding domains to FOXO1, FOXO4, NCOA1, or NCOA2 (Skapek et al., 2019). Given the abundance of transcriptional activation domains in the human proteome, the limited number of fusion partners suggests that routes to oncogenic transformation by transcriptional activation are quite constrained. Interestingly, all these fusion partners are activators that specifically interact with CBP/p300 rather than co-factors like NuA4 or BAF (Gerritsen et al., 1997; Lecoq et al., 2017; Nasrin et al., 2000). Our finding that ectopic SRF-C3orf62 expression preferentially affects SRF/MRTF targets over SRF/TCF targets suggests that co-factor/core promoter coupling (Haberle et al., 2019) could be involved in driving specific oncogenic gene expression programs. This could also explain why some but not other transactivation domains are frequently found in oncogenic fusions with DNA-binding transcription factors. Understanding how oncogenic fusion proteins are functionally connected to diverse co-activator complexes could provide therapeutic targets for emerging modulators of co-activator function (Brien et al., 2018; Vannam et al., 2021), paving the way to tailored therapy.

## ACKNOWLEDGEMENTS

We would like to thank Dr. Ben Piette for help with mass spectrometry analysis, Chris Mogg and Akashdeep Dhillon for help with experiments, Dr. Maria Vartiainen for SRF reporter constructs, and Drs. Tim Hughes, Michael Wilson, Maria Vartiainen, and Jussi Taipale for advice during the project. This project was supported by University of Toronto and Donnelly Centre startup funds, Canada Foundation for Innovation John R. Evans Leaders Fund grant and CIHR Project Grant (PJT175277) to M.T., and CIHR Foundation Grant (FDN143301) to A.-C.G. Proteomics work was performed at the Network Biology Collaborative Centre at the Lunenfeld-Tanenbaum Research Institute, a facility supported by Canada Foundation for Innovation funding, by the Government of Ontario and by Genome Canada and Ontario Genomics (OGI-139).

## AUTHOR CONTRIBUTIONS

Conceptualization: N.A., A.-C.G., and M.T.; Investigation: N.A., Z.-Y.L., A.-C.G., M.T.; Resources: Z.-Y.L.; Formal Analysis: N.A., A.-C.G., M.T.; Visualization: N.A., M.T.; Writing – Original Draft: N.A. and M.T.; Writing – Review & Editing: N.A., A.-C.G., and M.T, Supervision: A.-C.G. and M.T.; Funding Acquisition: A.-C.G. and M.T.

## DATA AVAILABILITY

The raw mass spectrometry data, alongside the annotation and identification files were deposited in the ProteomeXchange server MassIVE (massive.ucsd.edu) and assigned accession numbers MSV000087413/PXD025988 (AP-MS) and MSV000087424/PXD026006 (BioID). They can be accessed prior to publication at ftp://MSV000087413@massive.ucsd.edu (AP-MS; password: transcription) and ftp://MSV000087424@massive.ucsd.edu (BioID; password: transcription). All raw sequencing data is available at NCBI SRA under the accession number PRJNA729585. All reagents are available from the authors upon request.

## SUPPLEMENTARY FIGURES

**Figure S1.**
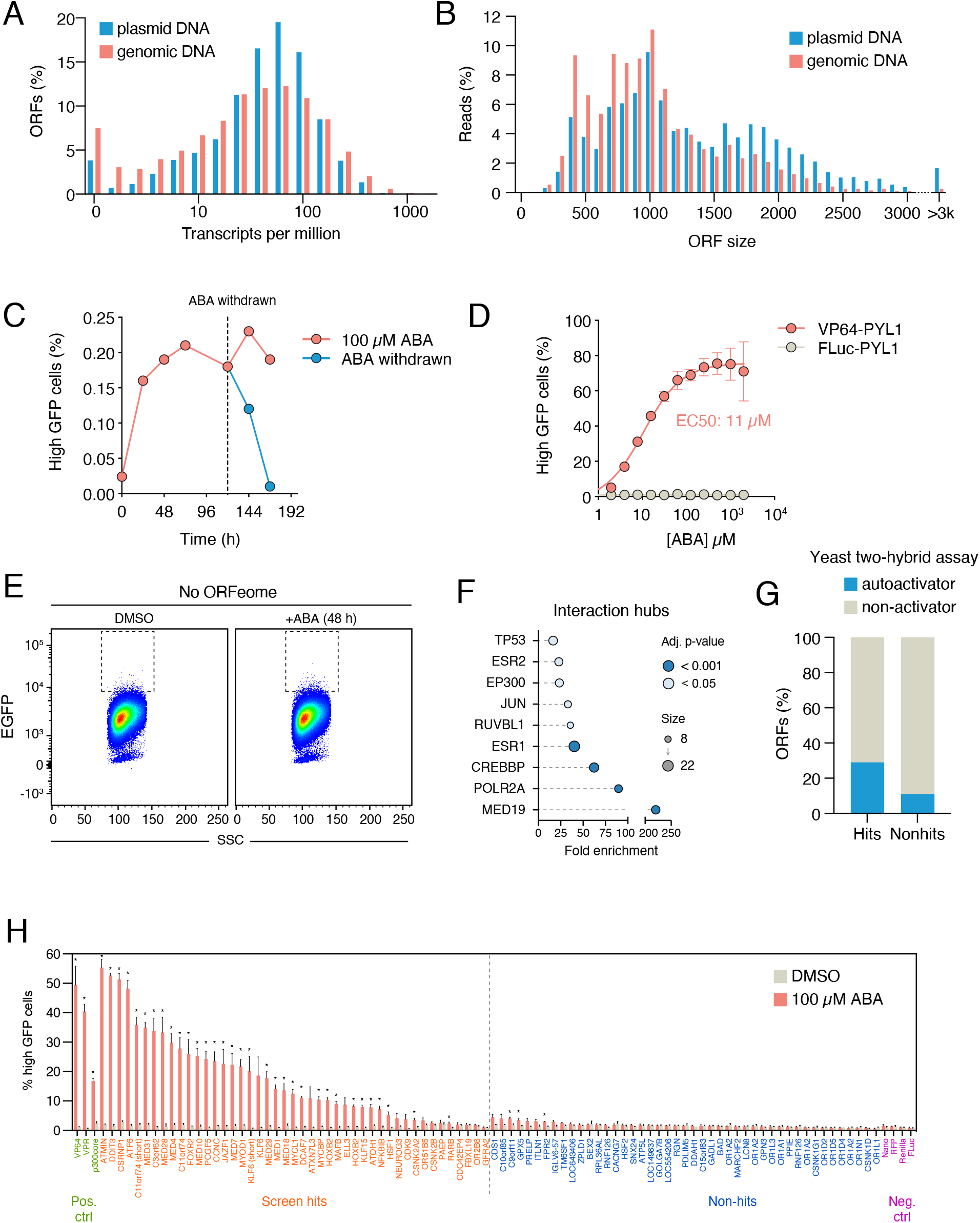
ORFeome screen for transcriptional activators. Related to Figure 1. **A,** Distribution of sequencing reads across the ORFeome in pooled plasmid DNA (blue) and in infected cells (red). **B,** Distribution of ORF sizes in the plasmid pool (blue) and in infected cells (red). **C,** Transcriptional activity depends on abscisic acid treatment. ORFeome-PYL1 infected cells were treated with 100 µM ABA and the fraction of high GFP cells measured by flow cytometer over time. **D,** The effect of ABA concentration on transcriptional activity. Reporter cells transfected with the indicated constructs were treated with increasing amounts of ABA for 48 hours. **E,** No high GFP population is observed in ABA treated cells not expressing the ORFeome-PYL1 library**. F,** Enrichment of interaction hubs among the hits of the activation screen. **G,** Enrichment of yeast two-hybrid autoactivators among the hits of the activation screen. **H,** Individual validation of transcriptional activators identified in the activation screen. Indicated constructs were transfected into the reporter cell line and high GFP cell fraction was measured by flow cytometry after 48-hour treatment with ABA. Asterisks indicate statistically significant ABA-dependent increase in high GFP population (FDR < 5%). Statistical significance was calculated with unpaired two-tailed t-test assuming equal variance, and corrected for multiple hypotheses with False Discovery Rate (FDR) approach of Benjamini, Krieger and Yekutieli (Benjamini et al., 2006).

**Figure S2.**
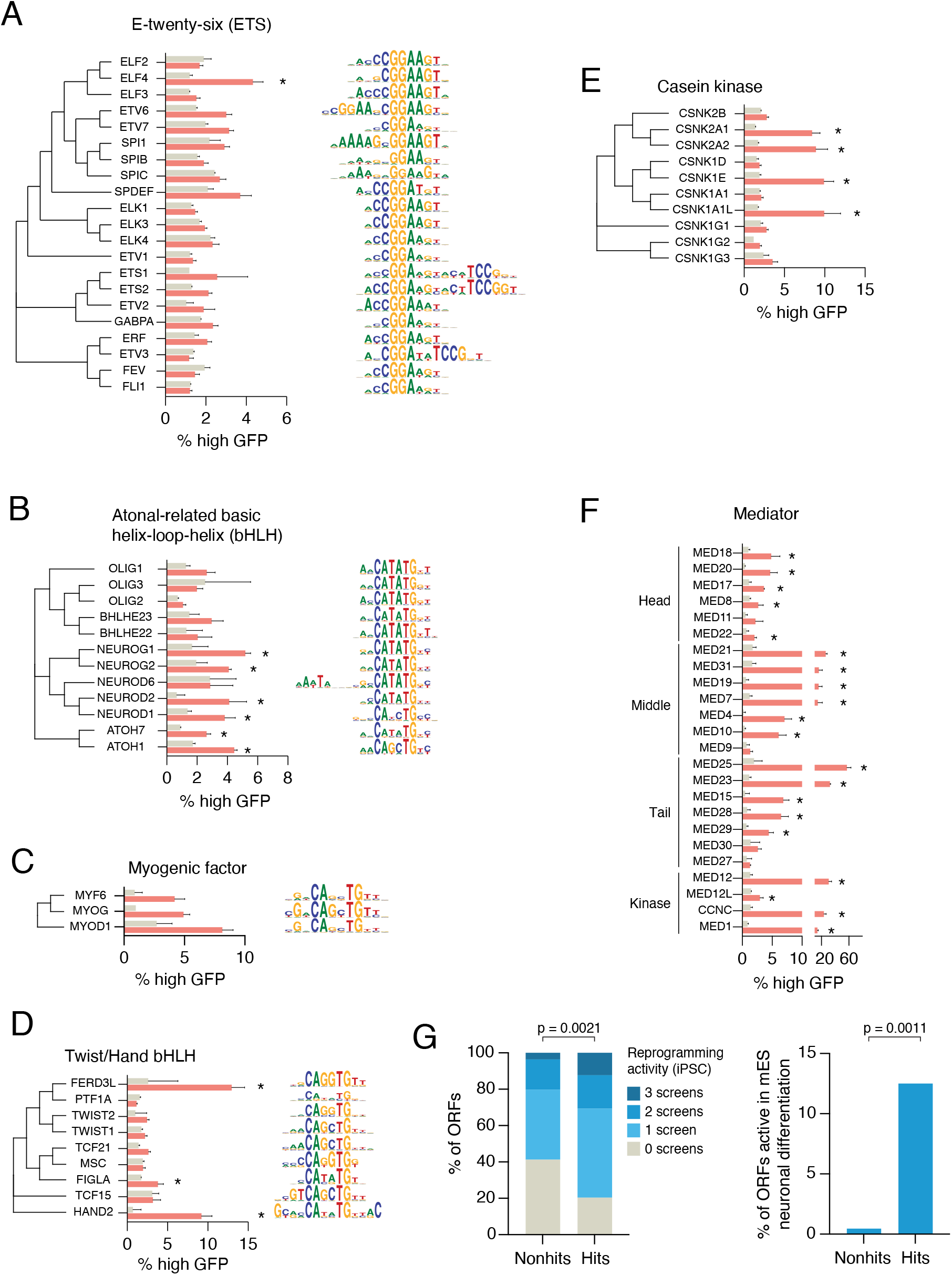
Transcriptional activity of transcription factor and chromatin-associated protein families. Related to Figure 2. Transcriptional regulators were individually tested for activation of the reporter in an arrayed manner. DNA-binding specificity (shown as sequence logos) is from CisBP (Weirauch et al., 2014). Asterisks indicate statistically significant activators (FDR < 5%). Statistical significance was calculated with unpaired two-tailed t-test assuming equal variance, and corrected for multiple hypotheses with False Discovery Rate (FDR) approach of Benjamini, Krieger and Yekutieli (Benjamini et al., 2006). **A,** ETS family TFs **B,** Atonal-related bHLH factors, **C,** Myogenic factors, **D,** Twist/Hand bHLH factors, **E,** Casein kinases, **F,** Mediator components, **G,** Transcription factors identified in the primary activation screen are enriched for factors that can reprogram human iPS cells (right) or mouse ES cells (left) when ectopically expressed (Ng et al., 2021; Theodorou et al., 2009). Statistical significance was calculated with a Wilcoxon rank sum test (left) or Fisher’s exact test (right).

**Figure S3.**
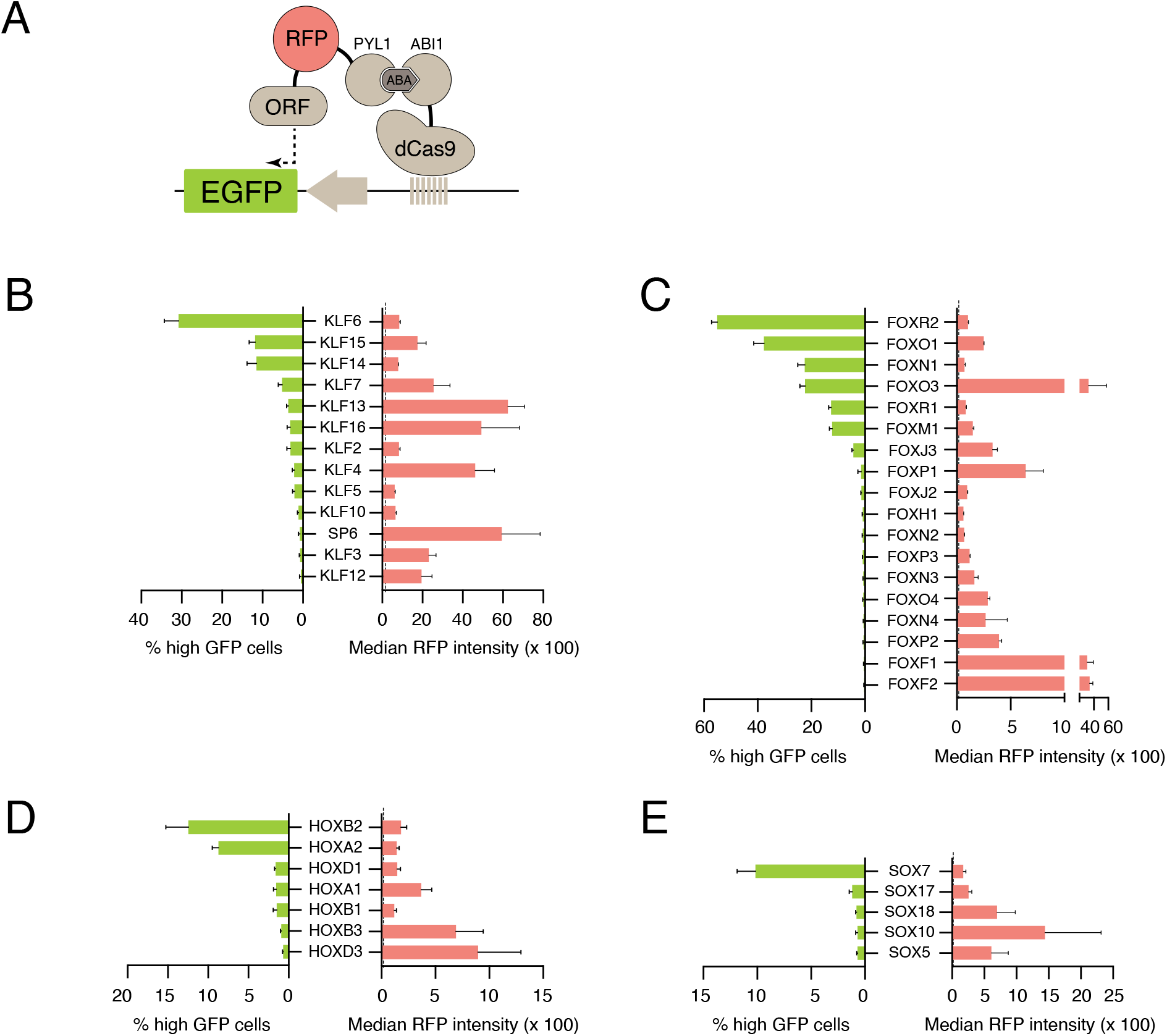
Differential activity of transcription factors is not explained by expression levels. Related to Figure 2. **A,** Schematic of the transcriptional activation assay measuring both reporter gene expression and effector protein expression. **B,** Transactivation of Kruppel-like factors (left) compared to the expression level of each factor as measured by RFP fluorescence (right). Background RFP intensity is shown as a dashed line. **C,** Forkhead TF activity. **D,** Homeodomain TF activity, **E,** SOX TF activity.

**Figure S4.**
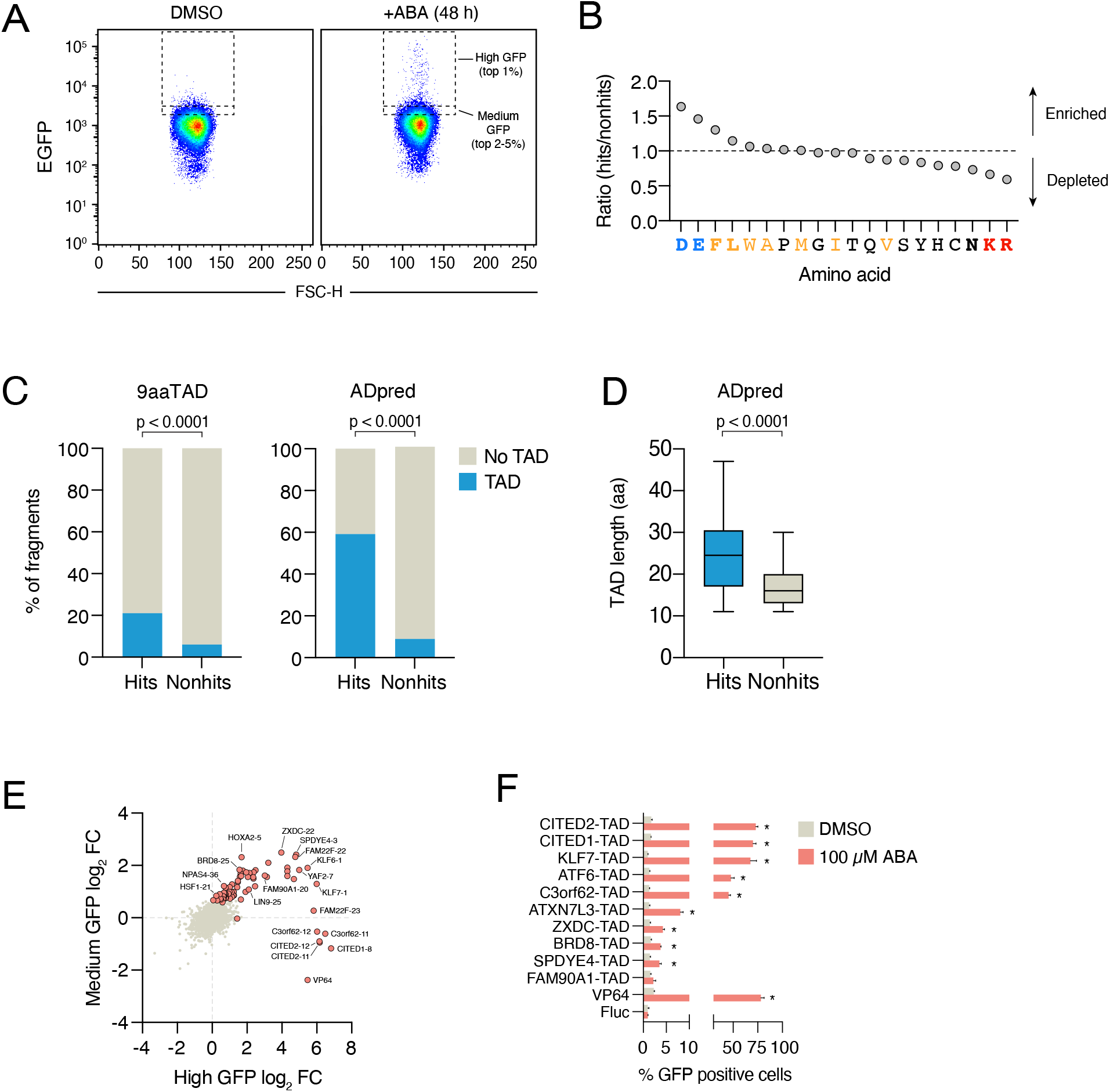
TAD-seq identifies transactivation domains. Related to Figure 3. **A,** High and medium GFP population was assessed by flow cytometry after recruiting 60aa fragments to the reporter with ABA for 48 hours. High GFP and medium GFP cells were sorted by FACS and ORFs enriched in the pools were identified by next-generation sequencing. **B,** Enrichment and depletion of amino acids among the identified transactivator fragments compared to inactive fragments in the library. Amino acids shown in bold were statistically significantly enriched or depleted. **C,** Enrichment of predicted transactivation domains among the active fragments. 9aaTADs were predicted with 9aaTAD prediction tool (https://www.med.muni.cz/9aaTAD/) using “Moderately Stringent Pattern”. Only 100% confident matches were considered in the analysis. ADpred algorithm was described in (Erijman et al., 2020). Statistical significance was calculated with Fisher’s exact test. **D,** Predicted TADs among active fragments are longer than those in inactive fragments. Statistical significance was calculated with two-tailed t-test assuming equal variance. **E,** Enrichment of fragments in the high GFP pool versus the medium GFP pool. Significant hits are shown as red circles. **F,** Individual validation of transactivating fragments. Indicated TADs were fused to PYL1 and transfected into the reporter cells in an arrayed format. Asterisks indicate statistically significant activators (FDR < 5%). Statistical significance was calculated with unpaired two-tailed t-test assuming equal variance, and corrected for multiple hypotheses with False Discovery Rate (FDR) approach of Benjamini, Krieger and Yekutieli (Benjamini et al., 2006).

**Figure S5.**
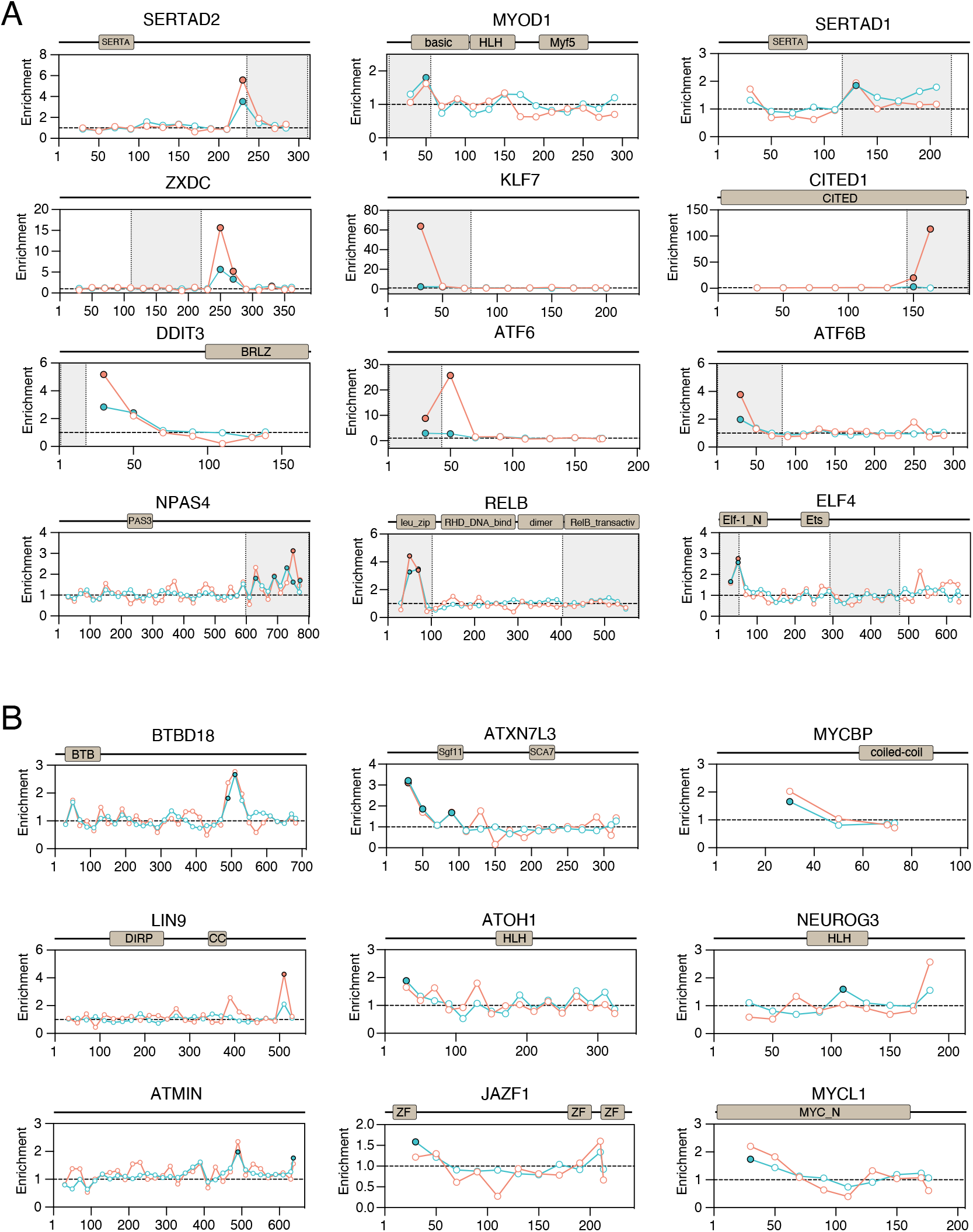
Systematic discovery of transactivation domains in human proteins with TAD-seq. Related to Figure 3. **A,** Examples of known transactivation domains. Domain organization is shown on top. TAD-seq plot shows the fold enrichment of RNAseq reads in the high GFP population (red) or the medium GFP population (blue). Each circle shows the mid-point (30th amino acid) of the 60-aa tile. Filled circles indicate statistically significant hits. Grey boxes indicate previously described transactivation domains. **C,** Examples of novel transactivation domains. Labeling is as in panel B.

**Figure S6.**
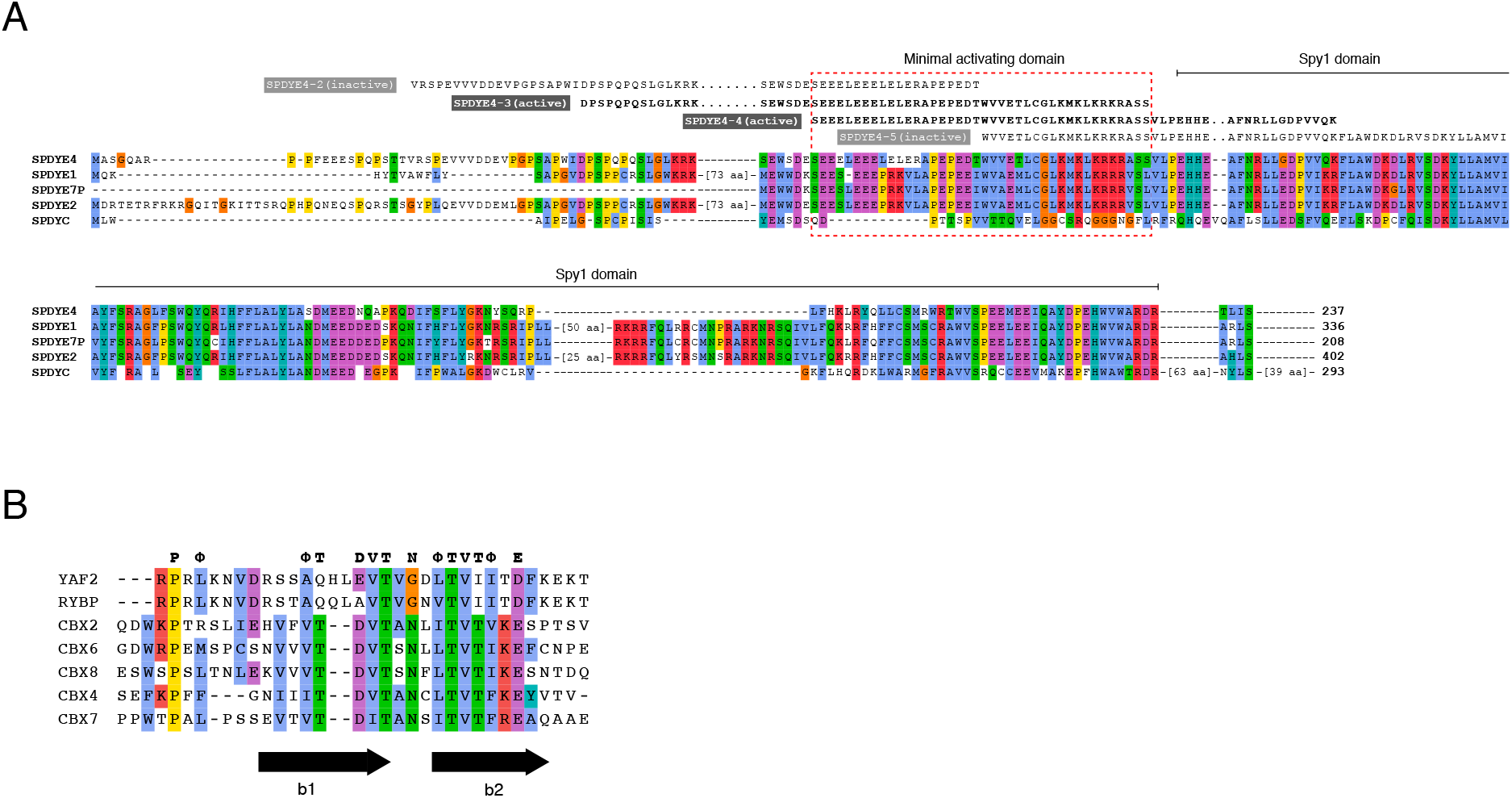
Transactivation domains of SPDYE4 and YAF2. Related to Figure 3. **A,** Alignment of five Spy1/RINGO family proteins. Four fragments screened by TAD-seq are shown above the alignment, and active fragments are indicated in bold. Red dashed box indicates the inferred minimal activating domain. Note that the region of the minimal activating domain is not conserved in SPDYC, which was the only Spy1/RINGO family member that did not activate transcription. **B,** Alignment of YAF2_RYBP domains of YAF2 and RYBP, and CBX_C domains of CBX family proteins. The location of the two beta sheets is indicated with arrows.

**Figure S7.**
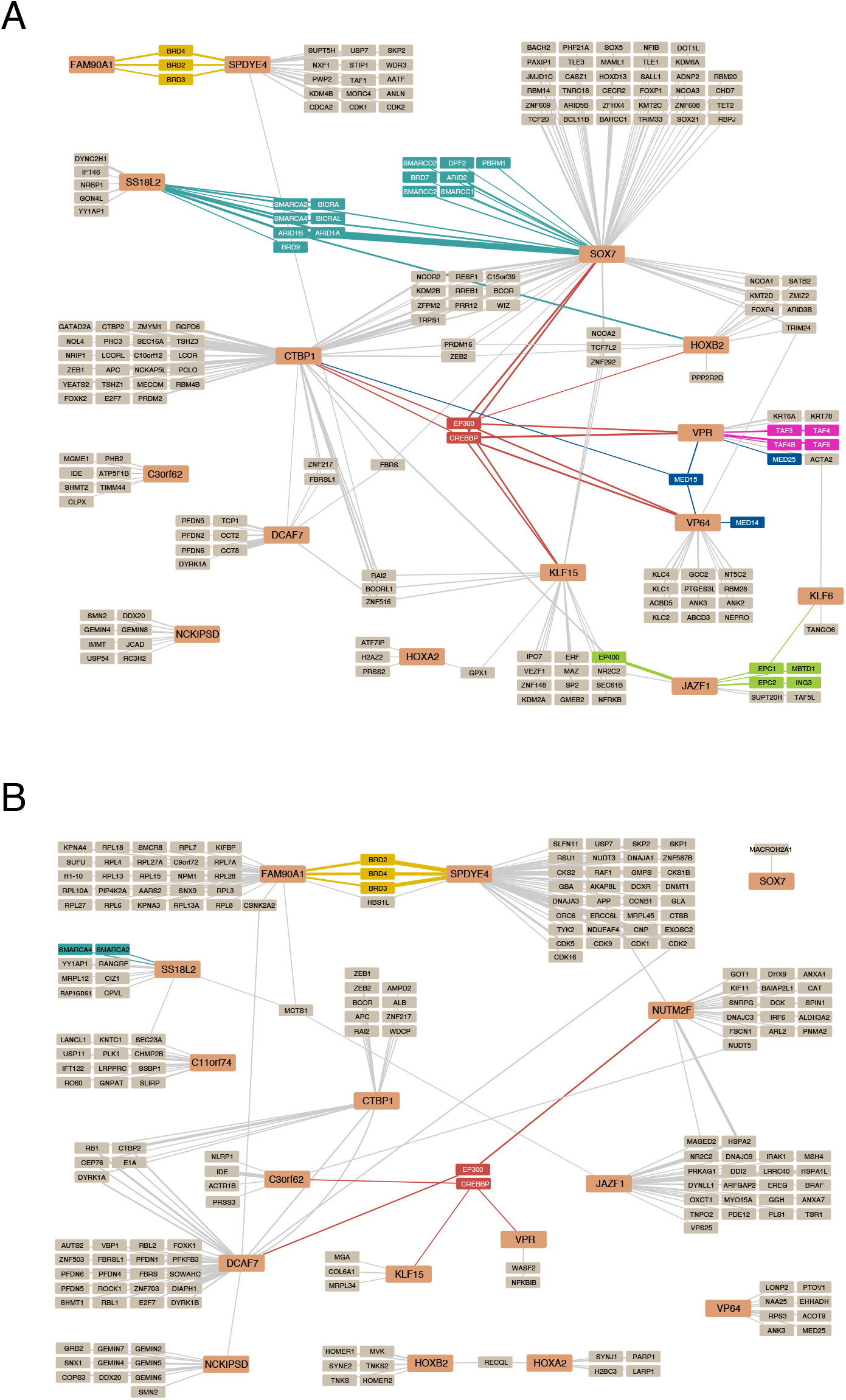
Interaction networks of transcriptional activators. Related to Figure 4. **A,** BioID2 network of transcriptional activators. Bait proteins are shown as light red rectangles. BAF complex members are shown in cyan, p300/CBP in red, NuA4 complex members in green, Mediator components in blue, and TFIID components in pink. The width of the edges indicates average spectral counts of two replicates. For clarity, two highly connected prey proteins (ZNF518A and ZNF518B) were removed from this visualization but are included in Table S1 **B,** AP-MS network of transcriptional activators. Labeling as in panel A. For clarity, nine highly connected prey proteins (APEH, ACTC1, ALDH1L1, PSDM4, LSS, FLII, PACSIN2, RPS3A, QPCTL) were removed from this visualization but are included in Table S1.

**Figure S8.**
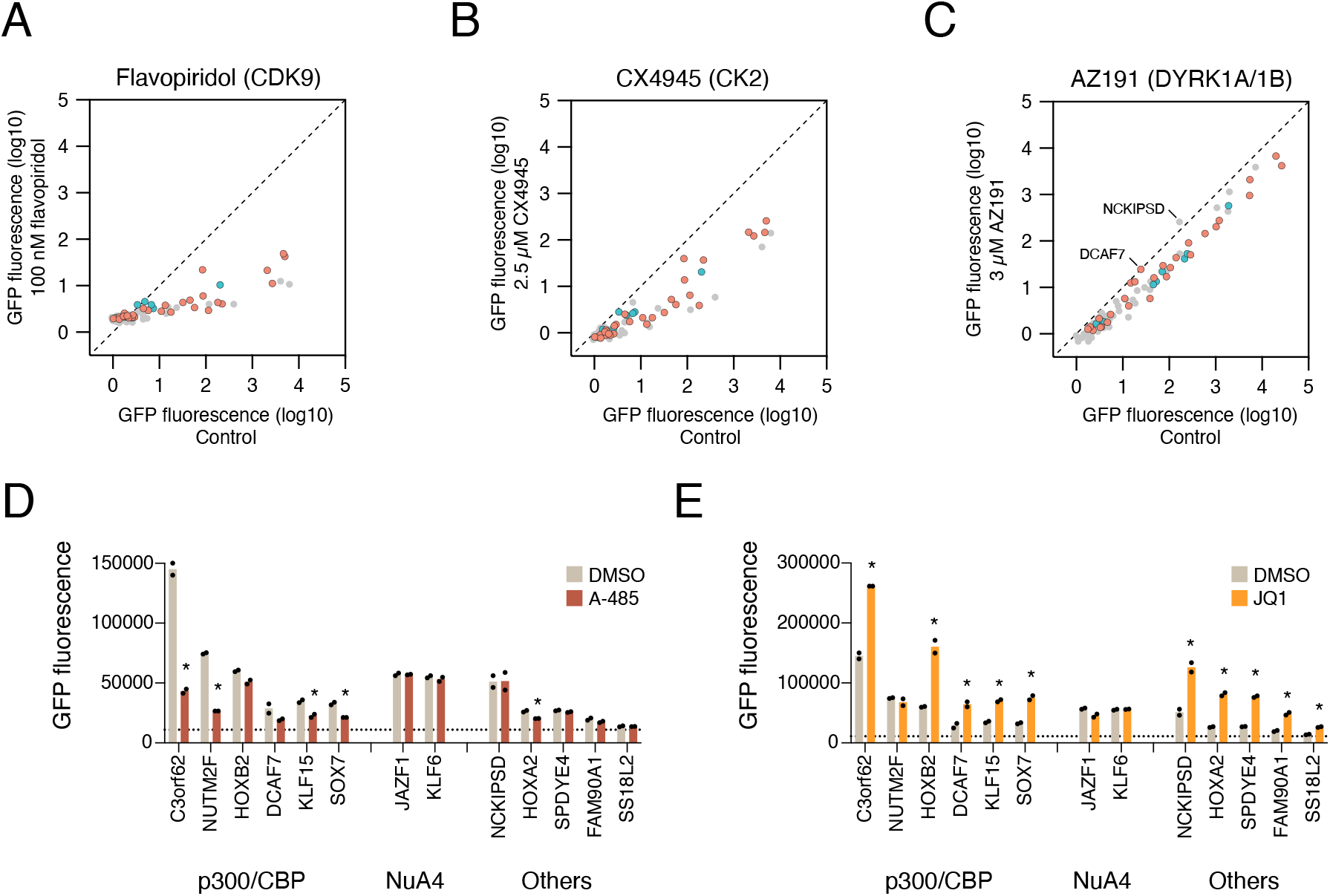
Effect of small molecule inhibitors on the transcriptional activity of 83 transcriptional regulators. Related to Figure 4. **A,** Effect of CDK9 inhibition by flavopiridol. Known p300 interactors are shown as red circles, known NuA4 interactors as blue circles. **B,** Effect of Casein kinase 2 inhibition by CX4945. Known p300 interactors are shown as red circles, known NuA4 interactors as blue circles. **C,** Effect of DYRK1A/DYRK1B inhibition by AZ191. Known p300 interactors are shown as red circles, known NuA4 interactors as blue circles. DYRK1A/DYRK1B interactor DCAF7, and DCAF7 interactor NCKISPD are highlighted. **D,** Effect of p300 inhibition by A-485 on transcriptional regulators characterized by AP-MS and BioID. Asterisks indicate statistically significant activators (FDR < 5%). Statistical significance was calculated with unpaired two-tailed t-test assuming equal variance, and corrected for multiple hypotheses with False Discovery Rate (FDR) approach of Benjamini, Krieger and Yekutieli (Benjamini et al., 2006). **E,** Effect of BET bromodomain inhibition by JQ1 on transcriptional regulators characterized by AP-MS and BioID. Asterisks indicate statistically significant activators (FDR < 5%). Statistical significance was calculated with unpaired two-tailed t-test assuming equal variance, and corrected for multiple hypotheses with False Discovery Rate (FDR) approach of Benjamini, Krieger and Yekutieli (Benjamini et al., 2006).

**Figure S9.**
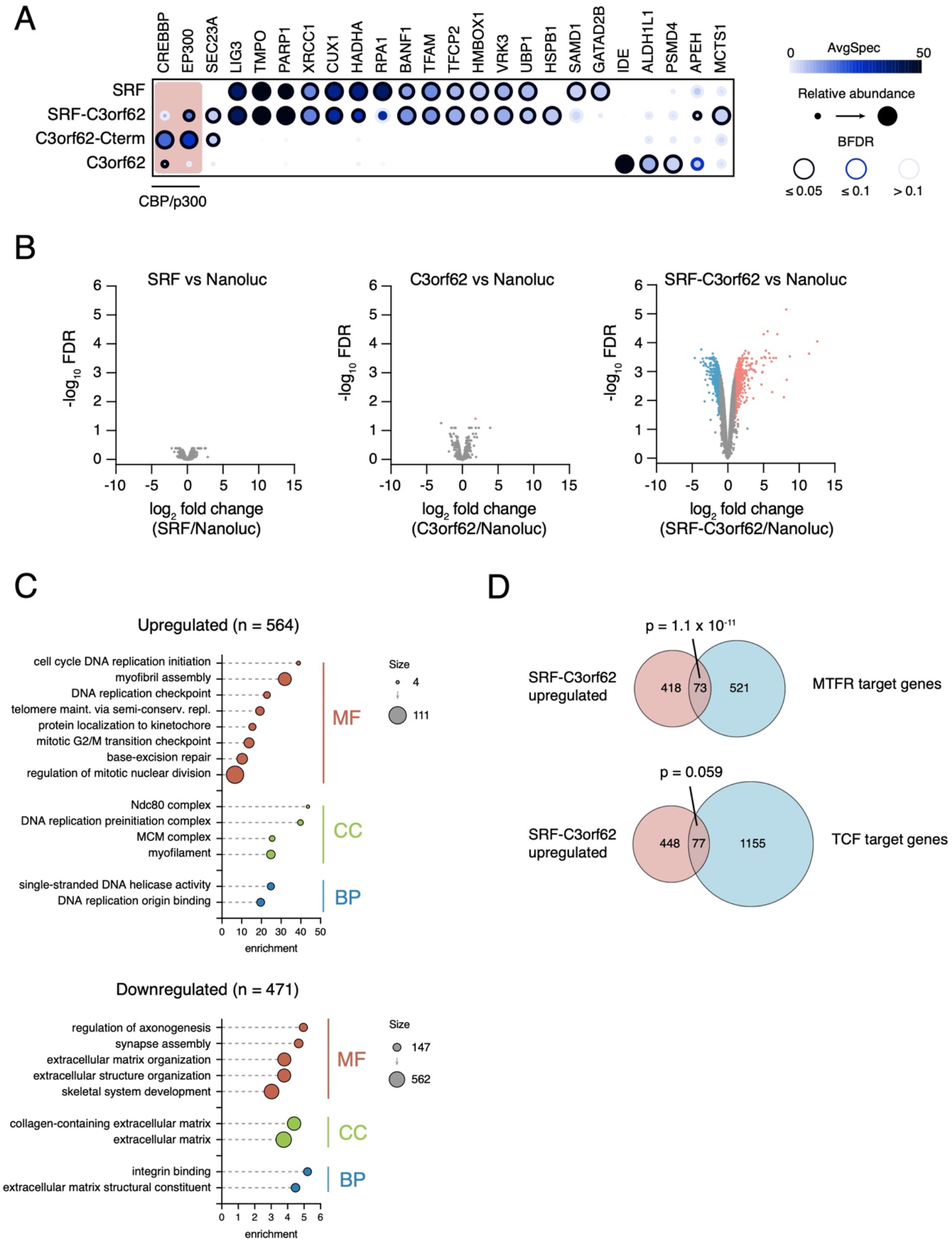
Transcriptional activation by SRF-C3orf62. Related to Figure 5. **A,** SRF, C3orf62, C3orf62-Cterm and SRF-C3orf62 interactors were characterized with AP-MS. SRF-C3orf62 fusion robustly interacts with CBP and p300. **B,** Analysis of differentially expressed genes in NIH3T3 cells expressing SRF-GFP, C3orf62-GFP, or SRF-C3orf62-GFP compared to cells expressing Nanoluc-GFP. Significantly upregulated (red) and downregulated (blue) genes are indicated. **C,** Gene Ontology enrichment analysis of significantly upregulated and downregulated genes in NIH3T3 cells expressing SRF-C3orf62-GFP. **D,** Overlap of genes significantly upregulatedy by SRF-C3orf62-GFP and target genes of SRF/MRTF or SRF/TCF, published previously (Esnault et al., 2014; Gualdrini et al., 2016). Statistical significance was calculated with a hypergeometric distribution test.

## METHODS

### Cell culture

All HEK293T cells, including the pTRE3G-EGFP reporter cell line (gift from Lei Stanley Qi lab, Stanford University) used for screens, were maintained in DMEM with 10% fetal bovine serum (FBS). NIH-3T3 cells were obtained from Dr. Sachdev Sidhu’s lab (University of Toronto) and maintained in Dulbecco’s Modified Eagle’s Medium (DMEM) with 10% bovine calf serum (BCS). All culture media were supplemented with 1% penicillin-streptomycin. Cells were maintained at 37°C in a humidified incubator at 5% CO2 and routinely tested for mycoplasma contamination.

### Lentivirus production

Lentiviral particles containing the pooled ORFeome and transactivation libraries were produced by transfecting 293T cells with pLX301-ORFs/TADs-PYL1, psPAX2 (Addgene #12260) and pVSV-G (Addgene #8454) at a ratio of 8:6:1. Transfection was performed using XtremeGENE 9 (Roche) on 15-cm dishes according to the manufacturer’s protocol. The medium was changed 6-8 hours post-transfection to harvest medium (DMEM + 1.1 g per 100 mL BSA). 72 hours after transfection, supernatant was filtered (0.45 μM), pooled and collected. A similar protocol was followed for small scale virus production when establishing individual stable cell lines with transfection being performed on 6-well plates using Lipofectamine 2000 (Thermo Fisher Scientific, 11668019) reagent.

### Cell line generation

A clonal line of the EGFP reporter line was generated expressing ABI-dCas9 (blastacidin, 6 µg/mL) and gRNA (co-expressing EBFP2) targeting the pTRE3G promoter. Single cells were sorted (FACS Aria IIIu, BD) and expanded and a clone showing induction by a strong transcriptional activator was selected for subsequent experiments. To generate NIH3T3 cells expressing doxycycline-inducible EGFP tagged proteins, entry cloned were picked from the hORFeome collection and subcloned into the Gateway compatible pSTV6-TetO-ccdB-EGFP lentiviral plasmid (a kind gift from Payman Samavarchi-Tehrani). NIH-3T3 cells were infected at the presence of 8 μg/mL polybrene and selected with 2 µg/mL puromycin 24 hours post infection.

### Pooled ORFeome library generation

Entry clones from the human ORFeome collection (v8.1) were collected into 40 standardized subpools each containing ∼384 ORFs and cloned into the lentiviral Gateway-compatible destination vector pLX301-DEST-PYL1. LR reactions were set up in duplicates with 150 ng of each entry ORF subpool, combined with 1 µl of Gateway LR clonase II in a total of 5 µl reaction volume and incubated overnight in TE buffer at room temperature. For the next two days, 1 µl additional LR enzyme was added in 4 µl TE and 150 ng destination vector to each reaction. Colonies were transformed into chemically competent DH5alpha *E. coli* and spread on LB agar plates containing carbenicillin (100 µg/µl) overnight at 30°C. Colonies were counted to ensure >200-fold coverage, collected in SOC on ice, pelleted and maxiprepped on multiple columns based on weight of the dry pellets.

### Activation domain tiling library generation

We generated a tiling library from 75 proteins identified as activators in the ORFeome screen. Oligonucleotides containing a 5’ adapter (GSQGSGSM) and a 3’ adapter (GGSVEREG) were synthesized as pooled libraries (Twist Biosciences). 6×50 µl PCR reactions were set up using NEBNext Ultra II Q5 master mix (New England Biolabs) with 5 nM oligos as template. PCR conditions were optimized to find the lowest cycle with a clean visible product at the expected 300 bp length. The thermocycling condition was an initial 30 s at 98°C, then 2 cycles of 98°C for 10 s, 63°C for 20 s, and 72°C for 15 s, followed by 10 more cycles of 98°C for 10 s and 72°C for 30 s with a final extension at 72°C for 5 min. Primers were designed to have Gateway compatible flanking sequences (Table S1). The resulting libraries were gel extracted by QIAgen gel extraction kit after loading on a 2% TAE gel for 2 hrs at 60V and subsequently cloned into pDONR221 using 20 separate BP reactions in total of 5 µl reactions. The entry plasmid pool was transformed after an overnight reaction into DH5alpha competent *E. coli* and incubated overnight on LB agar plates containing kanamycin (100 µg/µl). Colonies were collected and plasmid DNA purified. 20 LR reactions were then set up as described in the previous section. Each reaction was transformed into NEB 10-beta chemically competent *E. coli* and grown on LB agar plates containing carbenicillin (100 µg/µl) overnight at 30°C. Colonies were counted to ensure >200-fold coverage at each step of cloning, pooled and maxiprepped.

### Pooled activation screens

ORFeome and transactivation tiling libraries tagged at the C-terminus with PYL1 were packaged into lentiviral particles. A clonal EGFP reporter cell line stably co-expressing ABI-dCas9 and a gRNA targeting the promoter were transduced at low multiplicity of infection (MOI) with approximately 30% cell survival after puromycin (1 µg/mL) selection. Untransduced cells under the same condition were fully eliminated. Sufficient cells were transduced to maintain >500 fold coverage of the libraries. Recruitment was induced by treating cells with 100 µM abscisic acid (ABA, Sigma) for 48 hours. In parallel, a control batch of cells were treated with equal total volume of DMSO. Cells were then washed in PBS, treated with dissociation buffer (1 mM EDTA, 10 mM KCl, 150 mM NaCl, 5mM sodium bicarbonate, 0.1% glucose) and resuspended in flow buffer (5 mM EDTA, 25 mM HEPES pH 7, 1% BSA, PBS). High GFP population for each library, top 1% for ORFeome and two bins of top 1% and the next 4% for the TAD screen, were sorted and their genomic DNA directly extracted using QIAmp DNA Blood Mini Kit (QIAGEN).

### ORFeome sequencing

Nested PCR was performed using all the purified genomic DNA from sorted populations or at least 5 µg of genomic DNA from presort populations. The target ORFeome region was amplified from genomic DNA using primers targeting the T7 promoter and PYL1. The product of this reaction was pooled for each sample and further amplified by primers targeting outside the Gateway attB sites for an additional 10 cycles (Table S1). Amplicons were subsequently separated on 1% agarose gel and any visible PCR product excluding primer dimers were gel purified. After quantifying DNA using the Quant-iT 1X dsDNA HS kit (Thermo Fisher Scientific, Q33232), 50 ng per sample was processed using the Illumina DNA Prep, (M) Tagmentation kit (Illumina, 20018705), with 6 cycles of amplification. 2 µl of each purified final library was run on an Agilent TapeStation HS D1000 ScreenTape (Agilent Technologies, 5067-5584). The libraries were quantified using the Quant-iT 1X dsDNA HS kit (Thermo Fisher Scientific, Q33232) and pooled at equimolar ratios after size-adjustment. The final pool was quantified using NEBNext Library Quant Kit for Illumina (New England Biolabs, E7630L) and paired-end sequenced on an Illumina MiSeq.

### TAD sequencing

Performed nested PCR on the purified genomic DNA using primers targeting T7 promoter and PYL1 of the backbone vector in the first step creating a ∼470 bp product (Table S1). Products of the first reaction were then pooled and amplified for an additional 10 steps using primers targeting outside the Gateway sites (Table S1) creating a ∼300 bp product. Libraries were quantified on Qubit dsDNA Broad Range kit and paired-end sequenced on an Illumina MIseq with a custom PAGE-purified R1 sequencing primer (Table S1).

### Analysis of sequencing data from pooled activation screens

An index of the ORFeome reference sequences was created using the STAR aligner v2.7.8a with the length of the pre-indexing string set to 11 to account for the smaller ‘genome’ size. Reads from the ORFeome libraries were aligned with the STAR aligner allowing a maximum of 3 mismatches. For the TAD sequencing reads, cloning adapter sequences were first removed from both ends using cutadapt with CCAGTGTGGTGGAATTCTGCAGATATCAACAAGTTTGTACAAAAAAGTTGGCGGAAGTCAGGGTAGCGGAAGT for 5’ and CCGCCACTGTGCTGGATATCAACCACTTTGTACAAGAAAGTTGGGTAGCCTTCGCGTTCAACACTACCTCC for 3’ adapters. Bowtie reference was generated, and reads were mapped using Bowtie v1.2.3 allowing 0 mismatches. To identify activators, the edgeR package (Robinson et al., 2010) was used to calculate log2 fold change, p-value, and false discovery rate (FDR) for each ORF by comparing changes in counts from sorted samples to unsorted cells.

### Arrayed recruitment assays

Reporter cells stably co-expressing ABI-dCas9 and TetO gRNA were seeded either on 48-well or 96-well plates (Sarstedt) to reach 50-70% confluency on the day of transfection. 150 ng of each construct to be tested was transfected with polyethylenimine (PEI) at a ratio of 0.6 µl reagent. The day after transfection, recruitment was induced by treatment with ABA (100 µM). For tethered reporter assays in the presence of inhibitors, recruitment was similarly induced with 100 µM ABA the day after transfection but in the presence of either an inhibitor or the same volume of any additional DMSO. Inhibitors were dissolved in DMSO to a stock concentration of 10 mM. The final concentrations used were 100 nM for flavopiridol, 300 nM for JQ1, 1 µM for A-485, 2.5 µM for CX-4945 and 3 µM for AZ191. All inhibitors were a kind gift from the Structural Genomics Consortium (SGC). 48 hours after induction, cells were dissociated and resuspended in flow buffer using a liquid handing robot (TECAN) and analyzed by LSR Fortessa (BD). Cells were gated on high EBFP2 and RFP as a measure of gRNA and transfection control, respectively. Flow cytometry data was analyzed using FlowJo (v10).

### SRF reporter assay

30,000 NIH-3T3 cells on 24-well plates were transfected with 8ng SRF reporter (p3DA.luc), 20 ng reference reporter (pcDNA3.1-Nanoluc-3xFLAG-V5) and 50 ng 3xFLAG tagged constructs (Addgene #87063). Transfection was carried out using Lipofectamine 3000 reagent (Thermo Fisher Scientific, L3000001) according to the manufacturer’s protocols. Luciferase constructs were a kind gift from Dr. Maria Vartiainen (University of Helsinki). Cells were maintained in low-serum media (0.5% BCS) for 18 hours and stimulated for 7 hours (15% BCS), after which luciferase activity was measured. Firefly luciferase was normalized to Renilla luciferase activity using data from four independent transfections.

### RNA sequencing and analysis

NIH-3T3 cells with stable integrations of SRF, C3orf62, SRF-C3orf62 or Nanoluc tagged at the C-terminus with EGFP were induced with 1 µg/mL doxycycline for 24 hours. RNA was extracted from cells maintained in low-serum conditions (0.5% calf serum) for 22 hours using RNeasy purification kit (Qiagen) and treated with DNase on column. Samples were induced and collected in technical duplicates from 6-well plates. Libraries were prepared using the NEBNext Ultra II Directional RNA-seq with Poly-A selection kit, pooled and sequenced on a 100-cycle NovaSeq 6000 SP. Reads were aligned to the Gencode mouse primary assembly (GRCm39) with STAR aligner v2.7.8a. Counts for each gene were generated using the Gencode vM26 transcript annotations. Changes in gene expression compared to cells expressing Nanoluc-EGFP were quantified using the edgeR package (Robinson et al., 2010).

### Mass spectrometry samples

Entry clones were from the human ORFeome collection (Yang et al., 2011). Clones were transferred into pDEST-pcDNA5 vector carrying a C-terminal BioID2-FLAG tag (Kim et al., 2016) using Gateway recombinase. Stable HEK293 Flp-In T-REx cell lines were generated as previously reported (Piette et al., 2021).

For AP-MS, cells were grown to 70% confluence on 150 mm dishes before inducing bait expression with 1 µg/mL tetracycline for 24 hours. Cells were then washed once with 1xPBS, scraped, pelleted, flash-frozen, and stored at −80°C until processing. AP-MS was performed as previously described (Lambert et al., 2015). Briefly, cells were resuspended in cold lysis buffer (50 mM HEPES-NaOH pH 8.0, 100 mM KCl, 2 mM EDTA, 0.1% NP40, 10% glycerol, 1 mM PMSF, 1 mM DTT, 15 nM Calyculin A and protease inhibitor cocktail (Sigma-Aldrich P8340)) using a 1:4 pellet weight:volume ratio. Cells were lysed by one round of freeze-thaw, and lysates sonicated at 4°C using three 10-second bursts at 35% amplitude with 2 s pauses. Sonicated lysate was treated with 100U benzonase for 30 minutes at 4°C prior to clearing by centrifugation at 20,000g for 20 minutes at 4°C. An equal amount of supernatant from all samples processed within a batch was transferred to a tube containing 25 µL of pre-washed anti-FLAG magnetic bead 50% slurry (Sigma, M8823) and incubated for two hours at 4°C. Beads were recovered by magnetization and the supernatant discarded. Beads were washed once in lysis buffer, and once in 20 mM Tris-HCl pH8.0 with 2 mM CaCl_2_ and digested on beads with trypsin in two stages (1 μg trypsin for 4 hours followed by the addition of 0.5 μg trypsin to the supernatant and overnight incubation at 37°C), as previously described (Taipale et al., 2014). Finally, samples were acidified with 5% formic acid (final concentration) and stored at −80°C.

For BioID, cells were grown to 70% confluence in 150 mm dishes before inducing gene expression with 1 µg/mL tetracycline for 18 hours. 50 µM biotin was then added to each plate for 6 hours. Cell pellets were collected as for AP-MS, and resuspended in lysis buffer (50 mM Tris-HCl pH 7.5, 150 mM NaCl, 0.1% SDS, 1% Igepal CA-630, 1mM EDTA, 1 mM MgCl_2_, protease inhibitor cocktail (Sigma-Aldrich P8340, 1:500), and 0.5% sodium deoxycholate) using a 1:10 pellet weight:volume ratio. After sonication, each sample was treated with 250U Turbonuclease (BioVision 9207-50KU) and 1 µL RNase A solution (Sigma-Aldrich R6148) and incubated for 30 minutes at 4°C. SDS was then added to a final concentration of 0.25% and after mixing the samples were incubated for another 10 minutes at 4°C followed by centrifugation at 20,000g for 20 minutes. The supernatant was transferred to a tube containing 30 µl of pre-washed packed streptavidin beads (GE Healthcare, 17-5113-01). Streptavidin pulldown was done for 3 hours at 4°C. Beads were washed once in 1 ml of SDS buffer (2% SDS/50 mM Tris-HCl pH7.5), once in 1 ml lysis buffer, and once in TAP buffer (50 mM HEPES-KOH pH 8.0, 100 mM KCl, 10% glycerol, 2 mM EDTA, 0.1% Igepal CA-630), followed by three 1 ml washes with 50 mM ammonium bicarbonate pH 8.0. After the washes, beads were resuspended in ABC buffer containing 1 µg trypsin and incubated overnight at 37°C. The following day, the supernatant was collected and the streptavidin beads were washed with 50 µl water, which was combined with the first supernatant fraction. 0.5 µg trypsin was added to the combined supernatant sample, which was then incubated at 37°C for 4 hours. Beads were then spun down and the supernatant recovered. Beads were rinsed twice using ABC buffer and these rinses were combined with the original supernatant. Combined supernatants were dried by centrifugal evaporation.

### Mass spectrometry data acquisition and analysis

Samples were analyzed on a TripleTOF 5600 instrument (AB SCIEX, Concord, Ontario, Canada) as previously described (Piette et al., 2021) using Data-Dependent Acquisition (DDA). Data were processed and analyzed as previously described (Piette et al., 2021), using Proteowizard (Adusumilli and Mallick, 2017) implemented in ProHits v4.0 (Liu et al., 2016), Mascot, and Comet (Eng et al., 2013). The results were subsequently analyzed with the Trans-Proteomic Pipeline using iProphet (Shteynberg et al., 2011), and proteins with an iProphet probability ≥ 0.95 were further analyzed.

Significant interactors were identified with SAINTexpress (Teo et al., 2014). EGFP-BioID2-FLAG and EGFP-BioID2-FLAG were used as negative controls, using 2-fold compression for stringency as previously described (Mellacheruvu et al., 2013). SAINTexpress analysis used default parameters, and prey proteins were considered significant if they passed calculated Bayesian FDR cutoff of ≤5%. Dot plot figures were generated with ProHits-viz webserver (Knight et al., 2017).

